# Quantitative imaging of calcium dynamics with a green fluorescent biosensor and fluorescence lifetime imaging

**DOI:** 10.64898/2026.04.10.717680

**Authors:** Andrea Caldarola, Sebastián Palacios Martínez, Joachim Goedhart

## Abstract

Genetically encoded biosensors are GFP-based tools that can visualize the dynamics and spatial features of cellular processes. The design of a genetically encoded biosensor dictates the method that is used to measure the response. Common read-outs use some sort of fluorescence intensity measurement, which is subject to both technical and biological perturbations, including sample drift, excitation power fluctuations, changes in sample size/volume, or a change in expression level. Yet, the fluorescence lifetime of a fluorophore is not affected by the aforementioned perturbations. Therefore, biosensors that respond with a large lifetime change offer a more robust method of detecting cellular processes. Here, we report on protocols for calcium imaging using fluorescence lifetime imaging microscopy (FLIM) to measure the response of a genetically encoded lifetime biosensor. The protocols include details on biosensor production and purification, calibration of purified biosensor with FLIM, introduction of the plasmid in HeLa and endothelial cells, and timelapse analysis of FLIM data. In this chapter we use the green fluorescent biosensor G-Ca-FLITS as an example but the protocols can be generally applied to biosensors with lifetime contrast.

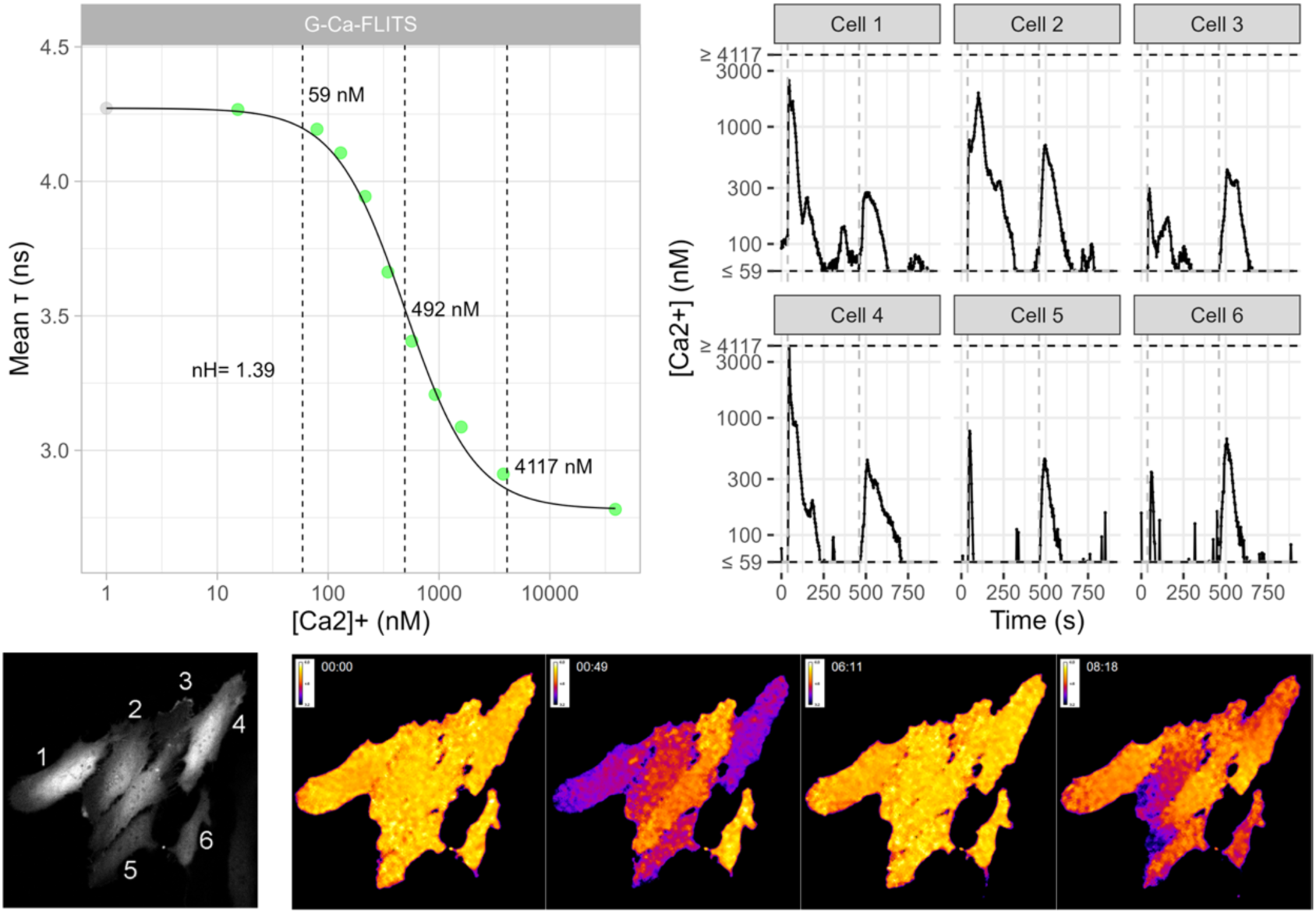

## 1. Introduction

Genetically encoded fluorescent biosensors use fluorescent proteins as building blocks. The biosensors can report on a large variety of analytes, co-factors and post-translational modifications. For the detection of the response, several read-outs are employed, including intensity change, FRET change, excitation ratio change, or re/translocation [1].

The read-out is dictated by the sensor design. Two popular designs that use a different read-out are the single fluorescent protein-based design and the FRET-based design [2]. Both designs use a “sensing” module, which is a protein domain that responds with a conformational change. The mechanism by which the the conformational change is translated into an optical signal is different for the two designs.

In the single fluorescence protein design, the conformational change in the sensing module perturbs the chromophore environment of the connected fluorescent protein [3]. As a result of the perturbation, the fluorescence intensity changes. The change in intensity is caused by a change in the extinction coefficient, a change in quantum yield, or both.

In the FRET based biosensor design, the conformational change changes the distance or orientation of a FRET pair. As a result, the FRET efficiency between the pair of proteins changes and the donor and emission intensity changes in opposite directions [4].

The classic read-outs of the single FP and FRET-based biosensors designs employ some sort of fluorescence intensity measurement. Yet, the fluorescence intensity is subject to both technical and biological perturbations, including, sample drift, excitation power fluctuations, changes in sample size/volume, or a change in expression level. In contrast, the fluorescence lifetime (a.k.a. excited state lifetime) is not affected by the aforementioned perturbations, offering a more robust way to read-out the sensor response [5, 6].

In theory, both sensor designs can also be analyzed using lifetime as the read-out. In reality, this option is hardly available. The single FP-based biosensors are usually based on GFP and the sole criterion in screening campaigns is a large intensity change. It turns out that almost exclusively, the resulting biosensors show a change in extinction coefficient, which does not affect the lifetime. There are a couple of GFP-based biosensors that show a significant lifetime contrast, i.e. HyPer3 [7], Peredox [8], sDarken [9], and qMaLioffG [10]. Intriguingly, all of these variants have a mutation at position 203, i.e. HyPer3; 203F, Peredox; 203I, sDarken; 203V, qMaLioffG; 203Y. Residue 203 is introduced in YFP to cause a red-shift [11], but it may well be that in GFP-based biosensors this residue is involved in generating lifetime contrast. Besides GFPs, also some single-FP biosensors that use a red fluorescent protein have lifetime contrast, i.e. RCaMP1h [12] and jRCaMP1b [13].

For FRET based biosensors, the change in FRET changes the quantum yield of the donor and this should be accompanied by a change in lifetime. In practice, most FRET based biosensors show hardly any lifetime change [14], with only a few exceptions [15, 16].

We reasoned that mTurquoise2 has great potential for generating single-FP based biosensors with lifetime contrast, since it has a high QY, and a low pKa [17]. So, we hypothesized that any intensity change would be due to QY changes as the (de)protonation is unlikely at physiological pH. Indeed, the GECI that we engineered using a circular permuted mTurquoise2, Tq-Ca-FLITS, shows a QY change and consequently a large lifetime change of ∼1.3 ns. In the meantime, other mTurquoise-based biosensors have been developed that show a lifetime change [18–20], suggesting that mTurquoise is a good template for lifetime biosensors. Because of the large lifetime change, the biosensor is well suited for quantitative imaging with FLIM. In fact, we were able to demonstrate its use in primary cells, organoids and in vivo [21].

Further engineering by introduction of the V27A, T81Y (which corresponds to the “YFP-mutation” T203Y), and N271D mutations (all in Tq-Ca-FLITS numbering) to the sensor resulted in the development of G-Ca-FLITS, which has a green-shifted spectrum similar to GFP and an increased lifetime change. G-Ca-FLITS maintains a high brightness in both the Ca^2+^ bound and unbound states, facilitating robust quantitative FLIM measurements [21].

Although the principles of quantitative imaging are applicable beyond imaging calcium, the number of biosensors with a good lifetime contrast is scarce. Since, in our hands, the variation in lifetime values because of technical reasons is ∼0.1 ns, we do not recommend using biosensors with a maximal lifetime change that is <0.3 ns. We, rather arbitrarily, define that a lifetime contrast exceeds 1 ns for ideal biosensors and 0.5 – 1 ns for usable biosensors, assuming a basal lifetime of 1 – 4 ns.

Here, we provide protocols for *in vitro* sensor calibration and for measuring Ca^2+^ in mammalian cells (both an immortalized, and a primary cell line) with FLIM-based, single FP biosensors. These protocols use G-Ca-FLITS as an example biosensor, but can be more broadly applied to biosensors that have lifetime contrast.

## 2. Materials

### 2.1 Protein production and purification

1. Shaking incubator (capable of holding 250 mL Erlenmeyer flasks, shaking at 200 rpm, with temperature control up to 37 °C).
2. Refrigerated centrifuge for 14 and 50mL tubes capable of centrifugation at 2,147 g (e.g., Eppendorf Centrifuge 5810 R).
3. Refrigerated ultracentrifuge capable of centrifugation at 40,000 g (e.g., Thermo Scientific Sorvall LYNX 6000).
4. Standard laboratory scale.
5. Tube rotator (e.g., Labinco LD-79) in an environment refrigerated at 4 °C.
6. HisPur™ Cobalt Resin(Thermofisher #89965) and Pierce Centrifuge Columns, 5 mL (Thermofisher #89897) or equivalent Cobalt/Nickel-based protein purification system.
7. Cytiva Disposable PD-10 Desalting Columns (Thermo Fisher # 11768488) or equivalent sample buffer exchange system.
8. Chemicals and Enzymes:

a. Lysozyme from chicken egg white (Sigma Aldrich #L6876).
b. Benzonase (Sigma Aldrich #E1014).
c. Halt™ Protease Inhibitor Cocktail (Thermo Fisher #87786).
9. Competent E. Cloni 10G cells (Sigma Aldrich #LGC601071) or equivalent high exogenous protein producer *E.coli* strain compatible with rhamnose induction.
10. Plasmid DNA encoding G-Ca-FLITS: pFPO-His-G-Ca-FLITS (Addgene #191455).
11. Buffers and Media, all diluted in MilliQ water unless otherwise stated:

a. Super Optimal Broth: 0.5% w/v Yeast Extract, 2% w/v Tryptone, 10 mM NaCl, 20 mM MgSO4, 2.5 mM KCl.
b. 100 mg/ml Kanamycin (Sigma Aldrich #K1377).
c. 40% w/v L-Rhamnose (Sigma Aldrich #W373011).
d. 10% NP-40 (United States Biological #N3500).
e. ST Buffer: 20 mM Tris, 200 mM NaCl, pH 8.
f. Equilibration buffer: 5 mM Imidazole in ST buffer.
g. Wash buffer: 15 mM Imidazole in ST buffer.
h. Elution buffer: 150 mM Imidazole in ST buffer.
i. Desalting buffer: 10 mM TrisHCl, pH 8.0.
12. Liquid Nitrogen.

### 2.2 *In vitro* calibration of the lifetime response

1. Calcium Calibration Buffer Kit #1 (Thermo Fisher #C3008MP).
2. Microscopy-grade glass bottom 96 well plates (e.g., Eppendorf #0030741030).
3. FLIM capable (confocal) microscope and FLIM imaging software. Protocols and analysis on this chapter use a time-domain FLIM Leica Stellaris confocal microscope with an environmental control chamber, on a computer with the LAS X FLIM/FCS software (Protocols were written for version 4.8.1).
4. Example data: DOI: <u>10.5281/zenodo.19357685</u>
5. Code: https://github.com/AndreaCaldarola/MMB_Quantitative_imaging_of_calcium_dynamics
6. Data processing and analysis software:

a. R and R-Compatible Integrated Development Environment (e.g., RStudio).
b. Python and Python-compatible Integrated Development Environment (e.g., JupyterLab or Spyder), preferably installed in a new Conda environment.
c. Microsoft Excel or similar spreadsheet editing software.

### 2.3 HeLa cell culture

1. Sterile cell culture hood.
2. Incubator with humidified chamber (37 °C, 7% CO_2_).
3. Dubbelco’s Modified Eagle Medium, GlutaMax Supplement (DMEM, Gibco #10566016), supplemented with 10% Fetal Bovine Serum (FBS, Gibco #A5256701).
4. Trypsin 0.05% (Gibco #25300062).
5. Wash buffer (HBSS or similar).
6. Vented-cap culture flasks (T25 or similar).

### 2.4 HUVEC cell culture

1. Sterile cell culture hood.
2. Incubator with a humidified chamber (37 °C, 5% CO_2_).
3. Endothelial Growth Medium 2 (EGM-2, PromoCell #C-22011), supplemented with 100 U/mL Penicillin and 100 μg/mL Streptomycin.
4. Phosphate Buffer Solution (Note 1).
5. Trypsin 0.05% (Gibco #25300062).
6. Trypsin neutralizing solution (TNS, Lonza #CC-5002).
7. 1 mg/mL Fibronectin (Sigma #F1141) dilute to a concentration of 10 µg/mL in PBS.
8. Cell culture containers (TC-treated vented-cap flasks, petri dishes or well plates) are all suitable).

### 2.5 PEI transfection of mammalian cell lines

1. Sterile cell culture hood.
2. Incubator with humidified chamber (37 °C, 7% CO_2_).
3. Mammalian cells at 50-60% confluency, plated on round coverglasses (24 mm, #1.5) in a 6-well plate, or on a glass-bottom 12/24-well plate (Note 2).
4. Cell culture medium (DMEM + 10% FBS).
5. Opti-MEM (or similar reduced-serum media).
6. PEI (1 mg/mL in sterile MiliQ water).
7. Plasmid DNA of interest (Note 3).

### 2.6 HUVEC electroporation

1. Sterile cell culture hood.
2. Incubator with humidified chamber (37 °C, 5% CO_2_).
3. HUVEC in culture (Note 4).
4. Endotoxin-free/low, high concentration plasmid DNA of interest (Note 5).
5. HUVEC culture reagents (See 2.4).
6. Neon Electroporation kit (Invitrogen #MPK10096, Note 6).

### 2.7 Timelapse FLIM of cells

1. FLIM capable (confocal) microscope. Protocols and analysis in this chapter use a time-domain FLIM Leica Stellaris8 confocal microscope with an environmental control chamber.
2. Mammalian cells expressing the biosensor of interest.
3. Low-fluorescence background imaging medium (Note 7).

### 2.8 FLIM Data analysis & conversion of FLIM data to [Ca^2+^] in live cells

1. Example data: DOI: <u>10.5281/zenodo.19357685</u>
2. Code: https://github.com/AndreaCaldarola/MMB_Quantitative_imaging_of_calcium_dynamics
3. Data processing and analysis software:

a. LAS X microscope software with FLIM and Phasor modules (Protocols were written for version 4.8.1).
b. R and R-Compatible Integrated Development Environment (e.g., RStudio).
c. Python and Python-compatible Integrated Development Environment (e.g., JupyterLab or Spyder), preferably installed in a new Conda environment.
d. Fiji distribution of ImageJ.
e. Microsoft Excel or similar spreadsheet editing software.

## 3. Methods

### 3.1 Protein production and purification

We use Thermofisher’s HisPur Cobalt Resin to bind and purify sensors expressed in bacteria transformed with inducible prokaryotic expression plasmids such as pDx (rhamnose induction) or pBAD (arabinose induction). In these plasmids a 6x-His tag is also added to the N-terminus of the sensor to enable its binding to the Cobalt ions present on the purification resin. We use the E. Cloni 10G strain for this purpose as they are compatible with protein expression-induction through rhamnose or arabinose as opposed to typical *E.coli* lab strains such as DH5α cells. Once we have produced and affinity-purified the sensor, we use PD-10 desalting columns to exchange the purification elution buffer with a storage buffer. The following protocol is specific to our experimental setup and thus we recommend thoroughly and carefully reviewing the user manual for the purification system you will use and making any necessary changes or adaptations.

Day 1:

1. Transform a vial of (Electrocompetent or Chemically competent) E.Cloni with pFPO-His-G-Ca-FLITS plasmid DNA .
2. After 1 hour of post-transformation recovery has passed, inoculate the transformed bacteria into 50 ml of Super Optimal Broth supplemented with 50 µg/mL Kanamycin and 0.4% w/v L-Rhamnose.
3. Grow the inoculated culture overnight at 37 °C shaking at 200 rpm in a 250 mL Erlenmeyer Flask.

Day 2:

4. Continue growing the bacterial culture shaking for 6 more hours at room temperature (Note 8).
5. Transfer the bacteria to a 50 mL falcon tube and centrifuge at 2147 g for 30 minutes at 4 °C to pellet the bacteria.
6. Discard the supernatant, resuspend the pellet by adding 20 mL ST buffer and gently pipetting up and down until the solution appears homogenous.
7. Centrifuge the tube at 2147 g for 30 minutes at 4 °C.
8. Discard the supernatant and resuspend the pellet by adding 5 mL ST buffer and gently pipetting up and now until the solution appears homogenous.
9. Freeze the bacteria at -20 °C (Notes 9, 10).

Day 3:

10. Fully defrost the frozen bacterial suspension on ice.
11. Add 5 mg of Lysozyme, 25 U of Benzonase and 50 μL Halt Protease Inhibitor, and incubate on ice for 30 minutes in order to lyse the cells, degrade the bacterial nucleic acids, and prevent degradation of the biosensor.
12. Meanwhile, prepare the protein purification beads:

a. Centrifuge 2 mL of Cobalt resin suspension for 2 minutes at 700 g in 14 mL tubes.
b. Discard the supernatant, resuspend the resin in 2 mL Equilibration buffer and once again centrifuge for 2 minutes at 700g.
c. Discard the supernatant, the beads are now ready for use.
13. Once the 30 minutes incubation is done, add 100 μL of 10% NP40 to the bacterial lysate, and centrifuge for 30 minutes at 40,062 g at 4°C (Note 11).
14. Transfer the visibly colored supernatant to the 14 mL tube with the purification beads, gently resuspending them in the process.
15. Add Equilibration Buffer until the tube is full, then rotate the tube for 30 minutes at 4 °C.
16. Centrifuge the tube at 700 g for 2 minutes.
17. Discard most of the supernatant, leaving ∼2 mL, then resuspend the beads and load the sample on a purification column.
18. Wash the beads with 2 mL Wash buffer 9 times, taking care to not let the beads completely dry out by adding the next Wash buffer volume before the last ∼1 mL of the previous has eluted.
19. Once the last washing step is completed, add 2 mL Elution buffer to the column to elute the purified sensors into Eppendorf tubes.
20.Wash a PD10 desalting column 5 times with 5 mL of Desalting buffer.
21. Load the purified sensor onto the column and elute with 3.5 mL of Desalting buffer (Note 12).
22. A small volume of the eluted sensor can be stored at 4 °C for short-term use (1-2 weeks). The rest can be aliquoted, snap-frozen in liquid nitrogen and kept at -80°C for long-term storage.

### 3.2 *In vitro* calibration of the lifetime response

We construct a calibration curve of the sensor’s lifetime response by imaging samples of G-Ca-FLITS at different free Ca^2+^ concentrations ([Ca^2+^]) prepared using Thermofisher’s Calcium Calibration Buffer Kit #1. We deem it necessary to use a buffering system (such as the one provided by the kit) due to the likely presence of small amounts of Ca^2+^ (which nonetheless render a calibration unreliable) in the tubes, pipette tips and well plates used to assemble and hold the calibration samples.

1. Dilute the purified sensor 1:100 in 10 mM K_2_EGTA and 10 mM CaEGTA buffer solutions.
2. Mix the 10 mM K_2_EGTA and 10 mM CaEGTA dilutions in different amounts in order to obtain samples with equal amounts of purified protein but different free [Ca^2+^], ranging from 0 to 39 μM (Notes 13, 14). The dilutions can be mixed directly within the wells of a glass-bottom 96 wells plate, with a final volume of 200 μL per sample (as in Table 1). The samples corresponding to the unmixed 10 mM K_2_EGTA dilution and the unmixed 10 mM CaEGTA dilution are referred to as the 0 [Ca^2+^] and the 39 μM [Ca^2+^] samples, respectively.
3. Image the calibration samples at a FLIM-capable microscope, in our case a LEICA Stellaris FALCON. Each well is imaged by acquiring a 512x512 image in its center using the same objective and imaging settings used during cell-based imaging of the sensor. In the case of G-Ca-FLITS this corresponds to a HC PL APO 40x/0.95 DRY objective, a 488 nm excitation laser line at 40 MHz frequency, and a HyDX detector with 495 - 575 nm as our detection range (Note 15). Pinhole size and frame accumulation can be increased as needed in order to detect >10^6^ photons for each sample, making sure to minimize any photon pile-up effects by staying under the 1 photon/laser pulse/pixel threshold and using low laser power (as in our cell-based imaging experiments, further explained in section 3.7).
4. Example data that can be opened in LAS X is available as “Calibration.lif” in the archive at zenodo (DOI: 10.5281/zenodo.19357685). In the LAS X software, ensure that each acquired image’s file name corresponds to the position of its well within the 96-wells plate (e.g. “G1”, “G2”, etc.). If necessary rename any of the acquired images by selecting the relevant file with the right mouse button and then selecting the “Rename” option in the drop-down menu that appears
5. In the LAS X FLIM window, select the acquired images by left clicking each file while holding down the Ctrl key.
6. Navigate to the Phasor Module of the LAS X FLIM window and once again select all images in the table displaying the data records of the loaded images in the bottom right of the software window (indicated as “H” in Fig. 6).
7. Right click the selected images and select the “Export Raw Data…” option from the drop-down menu that appears.
8. Type an appropriate name for the folder that will be created containing the raw photon counts of each image, such as “G-Ca-FLITS_Calibration”.
9. Thus you should obtain a folder such as “G-Ca-FLITS_Calibration.sptw” containing .ptu files named after the well that each sample was measured in, such as “G2.ptu”, “G3.ptu”, etc. The folder with exported data is available as “G-Ca-FLITS_Calibration.sptw” in the archive at zenodo (DOI: 10.5281/zenodo.19357685).
10. In Microsoft Excel, create a table with 3 columns containing the information of each sample, namely the sample name under a “Sample” column, the [Ca^2+^] of the sample under a “Ca” column, and the sensor’s name under the “Probe” column (as in Table 2).
11. Save the created table as a .csv file with an appropriate name such as “Calibration_Ca_Concentrations.csv”.

**Table 1.** Samples assembled in the example calibration dataset of G-Ca-FLITS, with corresponding free [Ca^2+^] at our imaging conditions (pH 7.2, 25 °C).

| Sample ID | G-Ca-FLITS diluted in 10mM K<sub>2</sub>EGTA Buffer (μL) | G-Ca-FLITS diluted in 10mM CaEGTA Buffer (μL) | Free [Ca<sup>2+</sup>] (nM) |
| --- | --- | --- | --- |
| G2 | 0 | 200 | 0.0 |
| G3 | 20 | 180 | 15.3 |
| G4 | 73 | 127 | 79.3 |
| G5 | 97 | 103 | 130 |
| G6 | 122 | 78 | 216 |
| G7 | 143 | 57 | 346 |
| G8 | 161 | 39 | 570 |
| G9 | 174 | 26 | 923 |
| G10 | 184 | 16 | 1587 |
| G11 | 193 | 7 | 3804 |
| F11 | 200 | 0 | 39000 |

**Table 2:** Table showcasing the correct format for the [Ca^2+^] dataframe.

| Sample | Ca | Probe |
| --- | --- | --- |
| G2 | 0.0 | G-Ca-FLITS |
| G3 | 15.3 | G-Ca-FLITS |
| G4 | 79.3 | G-Ca-FLITS |
| G5 | 130 | G-Ca-FLITS |
| G6 | 216 | G-Ca-FLITS |
| G7 | 346 | G-Ca-FLITS |
| G8 | 570 | G-Ca-FLITS |
| G9 | 923 | G-Ca-FLITS |
| G10 | 1587 | G-Ca-FLITS |
| G11 | 3804 | G-Ca-FLITS |
| F11 | 39000 | G-Ca-FLITS |

Once the raw photon counts of each sample have been exported as .ptu files, we use a Python and R-based data processing pipeline to generate a sensor calibration curve. The scripts and example calibration datasets relevant to this protocol can be found in the folder “3.2_In-vitro_fit-free_characterization_of_lifetime_response” of the github repository <u>MMB_Quantitative_imaging_of_calcium_dynamics</u>. In the “G_S_Calculator.ipynb” script, the binned photon counts of each sample are first converted into a 3-dimensional array using the python library ptufile [22]. In this array the first two axes correspond to the x and y axes of the image, while the z axis corresponds to a photon’s detection time. The first two axes are then summed together, obtaining a 1-dimensional array, in which to each binned photon arrival time corresponds a value which represents the amount of photons detected at that time. In the case of our imaging setup (images acquired with a pulsed laser with a frequency of 40 MHz and a HyDX detector), this corresponds to a 25.63 ns detection interval which is split into 264 bins, meaning each bin corresponds to a time interval of ∼0.097 ns (Note 16). Finally, using the FLIM_tools library [23], the average G and S coordinates of each sample are calculated.

12. Download the github repository <u>MMB_Quantitative_imaging_of_calcium_dynamics</u> as a zip, extract the files and move the folder “3.2_In-vitro_fit-free_characterization_of_lifetime_response” to an appropriate location on your computer.
13.Move the folder created in step 9 and the file created in step 11 into the “3.2_In-vitro_fit-free_characterization_of_lifetime_response” folder. In our sample dataset the latter 2 correspond to the “G-Ca-FLITS_Calibration.sptw” folder (available from zenodo, DOI: <u>10.5281/zenodo.19357685</u>) and the “Calibration_Ca_Concentrations.csv” file (included in the github repository).
14. In this folder, using the Anaconda Prompt or Anaconda Navigator, create a new conda environment with an appropriate name (e.g. “Sensor_Calibration”) and install the dependencies. For detailed instructions, see the Readme on the Github page: https://github.com/AndreaCaldarola/MMB_Quantitative_imaging_of_calcium_dynamics/tree/main/3.2_In-vitro_fit-free_characterization_of_lifetime_response
15. Start Jupyter Lab, and open the “G_S_Calculator.ipynb” file. This can be achieved by navigating to the folder containing the script file in Jupyter Lab’s file browser, accessed by clicking the folder icon on the left of the Jupyter Lab window or by holding the Ctrl, Shift and F keys on the keyboard.
16. If using a different dataset than the sample G-Ca-FLITS dataset provided, change the value for “in_dir” from ‘’G-Ca-FLITS_Calibration.sptw/’ to the name of the folder created on step 9.
17. In the top left of the Jupyter Lab window, select “Run” and then “Run All Cells” to execute the script.
18. Three new files should then be created in the folder containing the script: “all_decays.csv”, “all_decay_raw.csv” and “df_GS.csv”.

We then use the “Sensor Calibration.qmd” R script to characterize the sensor’s lifetime response to [Ca^2+^]. First, each sample is plotted in phasor space according to its G&S coordinates (Fig. 1A). The calibration continues by calculating, for each sample, the difference between its G and S and the G and S of the 0 [Ca^2+^] sample (i.e. its dG & dS), and then plotting them with a line connecting the 0 & 39 μM [Ca^2+^] samples (Fig. 1B). Each sample is projected onto the line and the distance of the projected data relative to the length of the line is calculated. This value is referred to a sample’s line fraction, which for each sample should result in a value between 0 (corresponding to the 0 [Ca^2+^] point) and 1 (corresponding to the 39 μM [Ca^2+^] point). The line fraction data is then fitted using non-linear regression via the R package nls.multstart [24], establishing the constants in the equation which dictates the relationship between the line fraction value of each sample and its [Ca^2+^] (Equation 1).

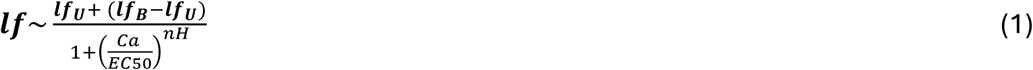

where *lf* is the line fraction value; 𝒍𝒇_𝑼_ is the minimum line fraction value, corresponding to the line fraction value of the 0 [Ca^2+^] sample in which G-Ca-FLITS’ Ca^2+^-unbound state is most abundant; 𝒍𝒇_𝑩_ is the maximum line fraction value, corresponding to the line fraction value of the 39 μM [Ca^2+^] sample in which G-Ca-FLITS’ Ca^2+^-bound state is most abundant; *Ca* is the free [Ca^2+^] that corresponds to the *lf* value; *EC50* is the [Ca^2+^] in which we determine the line fraction value to be the middle point in between the minimum and maximum line fraction values; and *nH* is the Hill coefficient of the sensor’s Ca^2+^-induced lifetime change.

Thus, we calculate the EC50 of G-Ca-FLITS’ Ca^2+^-induced lifetime change and its Hill coefficient. We then calculate the line fraction value for a range of [Ca^2+^] values using our equation parameters, and plot the resulting points together with the original calibration dataset in order to visualize the fitted equation (Fig. 1C). The calculated line fraction is then corrected for the intensity difference of the two states (Equation 2).

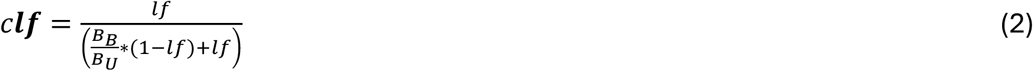

where *clf* is the corrected line fraction value; *lf* is the initial line fraction value, i.e. the apparent line fraction; 𝑩_𝑩_ is the molecular brightness of the Ca^2+^-bound state of G-Ca-FLITS; 𝑩_𝑼_: = is the molecular brightness of the Ca^2+^+-unbound state.

The corrected line fraction values represents the relative abundance of the sensor’s Ca^2+^-bound state which is present at a given [Ca^2+^], and can be fit and plotted in the same manner as the line fraction in order to determine G-Ca-FLITS’ true Kd (Fig. 1D).

19. In RStudio, open the “Sensor Calibration.qmd” file located within the “3.2_In-vitro_fit-free_characterization_of_lifetime_response” folder and install all package dependencies when prompted.
20. If not using the provided sample dataset, change the string that is used by “read.csv()” in the script, i.e changing “Calibration_Ca_Concentrations.csv” to the name of the file created on step 11.
21. Hold down the Ctrl, Alt and R keys (option+command+R on a Mac) on the keyboard in order to fully execute the script.
22. A new folder called “Plots” should be created, containing plots which showcase the results of the sensor characterization such as those shown in Fig. 1.

**Figure 1:**
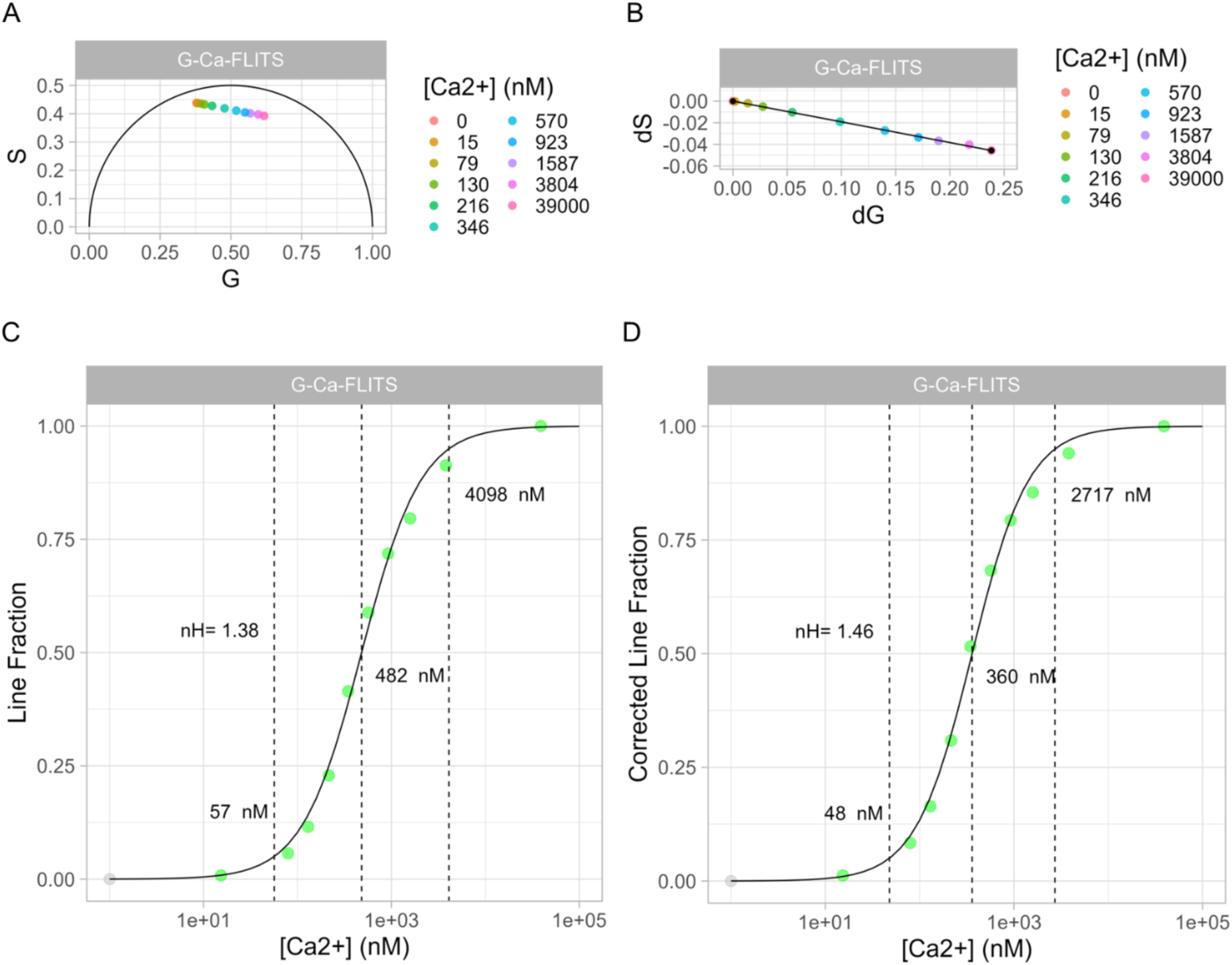
Results of the sensor characterization protocol applied to our sample G-Ca-FLITS calibration dataset. A) Phasor plot of the G-Ca-FLITS calibration samples: each sample’s average G & S are plotted. B) dG&dS plot of the G-Ca-FLITS calibration samples: each sample is plotted with coordinates corresponding to the difference between its G and S and the G and S of the 0 [Ca^2+^] sample. C) Fitted line fraction plot and D) Fitted corrected line fraction plot: each sample’s line fraction (C) or corrected line fraction (D) value is plotted as a function of its [Ca^2+^]. The 0 [Ca^2+^] sample (shown in gray) is plotted as 1 nM [Ca^2+^] in order to display it on a logarithmic axis. For both plots the Hill coefficient (nH), EC05, EC50 and EC95 values are shown, as well as a black line of points computed using the fitting results.

### 3.3 HeLa cell culture

We utilize a HeLa cell line (CCL-2) for our routine sensor development and characterization, due to their ease of culture and transfection. HeLa cells are cultured in DMEM + GlutaMax supplemented with 10% FBS. Cells are maintained at 37 °C, 7% CO_2_ and passaged twice per week at 1:10 or 1:5 dilution, depending on current and desired confluency.

1. For passaging, after removal of the old medium wash cells with a large volume (∼5 mL for a T25 culture flask) of wash buffer (HBSS).
2. Cover with trypsin and incubate at 37 °C until cells have fully detached (500-600 μL for a T25, 3-5 minutes).
3. Resuspend in DMEM (to 5 mL for a T25 flask) and pipette gently to break-up clumps of cells.
4. Pipette the required amount of cell suspension (e.g., 1 mL from a confluent flask will reach confluency again in 2-3 days) into a new flask with pre-warmed DMEM.

### 3.4 HUVEC cell culture

Human umbilical vein endothelial cells (HUVEC) can be kept in culture until passage 6-7 (close to two weeks culture time) before differentiating into fibroblast-like cells. Therefore, experiments that deal with endothelial biology should be conducted within this timeframe. Alternatively, other types of endothelial cells with longer culture times (e.g., BOEC) can be used. HUVEC are particularly sensitive to confluency, and should be cultured at a minimum confluency of around 30-50% (Note 17).

1. Warm up culture reagents to 37 °C and bring Trypsin solution to room temperature (Note 18).
2. Before passaging cells, add enough 10 μg/mL fibronectin so that the whole culture area is covered and incubate at 37 °C for at least 30 minutes. Following incubation, remove excess fibronectin before passaging cells onto the new surface.
3. Remove old medium from HUVEC to be passaged. Wash twice with 5 mL warm PBS (Note 1).
4. Add 10 μL trypsin/ 1 cm^2^ culture area (1 mL for a 100 mm petri dish is enough), incubate at 37 °C for 3 minutes.
5. Ensure cells have detached and add TNS at a 1:1 ratio to Trypsin. Gently mix the dish and rest for 30 seconds to neutralize Trypsin.
6. Resuspend in EGM-2 and passage cells at the desired confluency onto a fibronectin-coated culture surface with pre-warmed EGM-2.
7. HUVEC can survive over the weekend in the same medium, otherwise refresh the medium every other day (Table 3).

**Table 3:**
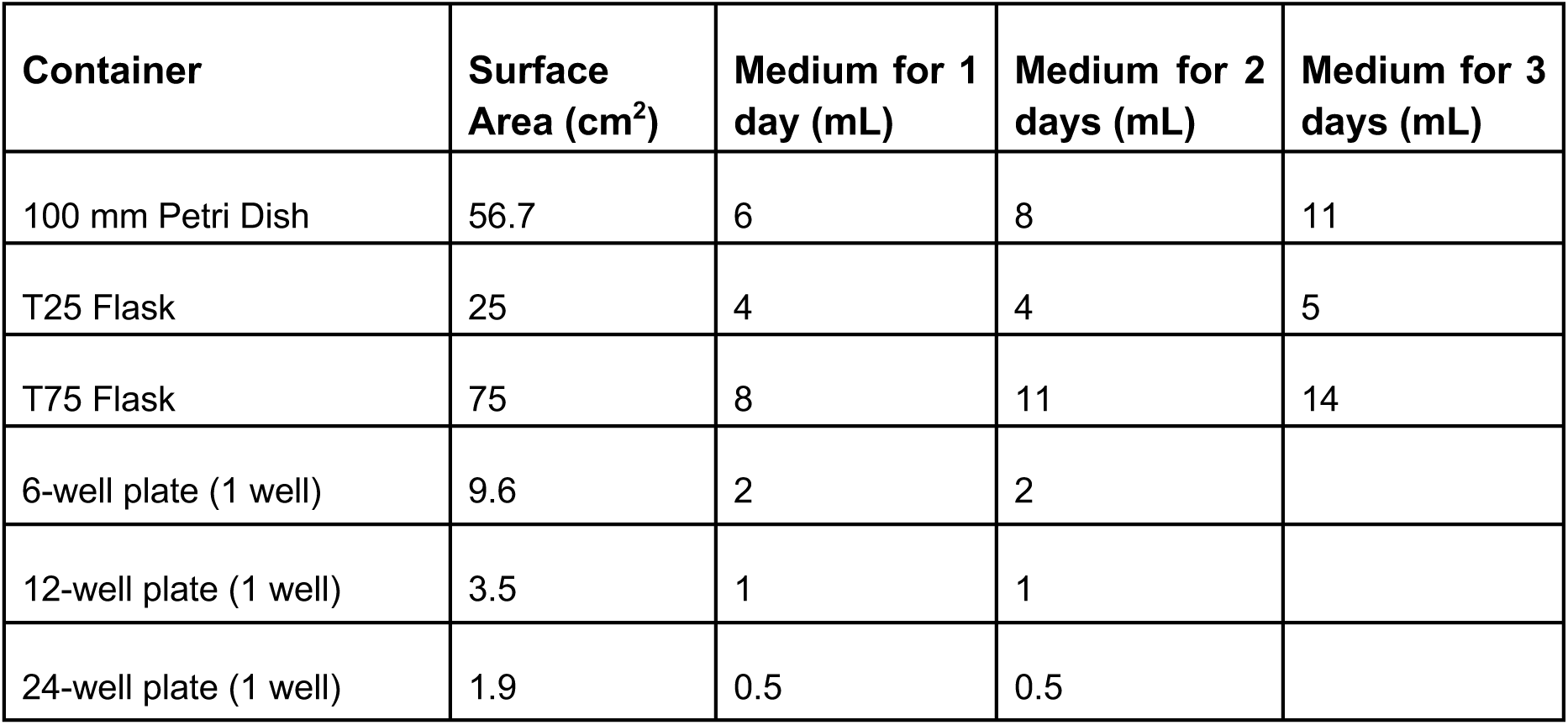
Suggested volumes of medium for culturing HUVEC.

| Container | Surface Area (cm<sup>2</sup>) | Medium for 1 day (mL) | Medium for 2 days (mL) | Medium for 3 days (mL) |
| --- | --- | --- | --- | --- |
| 100 mm Petri Dish | 56.7 | 6 | 8 | 11 |
| T25 Flask | 25 | 4 | 4 | 5 |
| T75 Flask | 75 | 8 | 11 | 14 |
| 6-well plate (1 well) | 9.6 | 2 | 2 |  |
| 12-well plate (1 well) | 3.5 | 1 | 1 |  |
| 24-well plate (1 well) | 1.9 | 0.5 | 0.5 |  |

### 3.5 PEI transfection of mammalian cells

We employ polymer carrier PEI transfection for delivery of plasmid DNA to standard mammalian cell lines (HeLa, U2OS, HEK, etc.). At the time of transfection, cells should be around 50-60% confluent.

1. 24 hours before transfection, plate cells by placing round coverglasses in a 6-well plate. For plating a full well plate from a confluent T25 flask, detach cells according to protocol 3.3 and bring 20% of total cell suspension to a final volume of 12 mL with DMEM + 10% FBS. Divide evenly (2 mL per well for 6 wells) and let cells attach overnight. Alternatively, cells can be plated directly onto glass-bottom, 12/24-well plates following the same steps.
2. Bring DNA, PEI, and Opti-MEM to room temperature.
3. For each transfection mix, we use a ratio of 1 μL PEI per 100 ng of plasmid DNA (Note 19), which is then diluted in 25-100 μL Opti-MEM (Table 4). Add the three reagents one at a time in a sterile 1.5 mL tube, adding the Opti-MEM last. Mix thoroughly by pipetting but avoid creating bubbles.
4. Incubate the transfection mix for 20 minutes at room temperature.
5. Add the transfection mix dropwise into the cell culture wells.
6. Incubate transfected cells at 37 °C, 7% CO_2_. Peak DNA expression occurs 24-48 hours post-transfection.

**Table 4:** Reagent volumes for PEI transfection calculated for different culture areas.

| Transfection area | DNA (ng)<br>(Note 19) | PEI (μL) | Opti-MEM (μL) | DMEM (mL) |
| --- | --- | --- | --- | --- |
| 6-well plate (per well) | 250 | 2.5 | 100 | 2 |
| 12-well plate (per well) | 125 | 1.25 | 50 | 1 |
| 24-well plate (per well) | 62.5 | 0.625 | 25 | 0.5 |
| Full 6/12/24-well plate | 1500 | 15 | 600 | 12 |

### 3.6 HUVEC electroporation

Commonly used lipid, or chemical transfection methods (PEI, lipofectamine, etc.) generally have a poor transfection efficiency for endothelial cells. We obtain highly efficient DNA delivery to HUVEC using electroporation, particularly with the Invitrogen Neon Electroporation system (Note 6).

1. Bring R and E2 buffers to room temperature. Pre-warm culture reagents.
2. Set the electroporator settings to 1300 V and 30 ns pulse duration (single pulse).
3. Fill one electroporation tube with 3 mL E2 buffer and place it in the electroporator (inside of the flow hood) with the electrodes facing each other ensuring it clicks in place.
4. Pipette 2 μg DNA of interest per transfection *n,* into a sterile 1.5 μL tube.
5. Detach HUVEC following cell culture protocol using trypsin and TNS, (See protocol 3.4). If necessary count cells, and spin cells down down at 200 G for 3 minutes.
6. Resuspend HUVEC in 120 μL R buffer per electroporation *n*.
7. Add 120 μL R buffer cell suspension into each 1.5 mL tube containing 2 μg DNA, mix gently by pipetting up and down. (Note 20).
8. Place a 100 μL electroporation tip on the electroporation pipette, check that the electrode is firmly gripped by the pipette and the plastic tip is correctly attached. Gently aspirate the cell and DNA suspension to the full volume of the tip, making sure that there are no bubbles in the liquid (Note 21).
9. Place electroporation pipette in the electroporator tube filled with E2 buffer and ensure it clicks into place. Press START on the control panel of the electroporator to deliver the electrical pulse and wait to hear a click from the machine. (Note 22).
10. If the electroporation is successful, gently pipette cells into a well pre-coated with fibronectin and containing pre-warmed EGM-2.
11. Let cells attach for 30 minutes to 1 hour and refresh medium (Note 23). Maximum expression occurs 24-48 hours post-electroporation, depending on construct size. HUVEC can be imaged in full EGM-2 medium.

### 3.7 Timelapse FLIM of live cells

The inherently (semi)-quantitative nature of FLIM biosensor imaging necessitates that extra care be taken to obtain robust measurements that can be accurately translated into [Ca^2+^]. A robust FLIM measurement can require between 100 and >100,000 photons, depending on the number of components, lifetime contrast, and detection window. It is also important to evaluate the fraction of photons which belong to the sensor itself, and those that can be categorized as “background”, including photons belonging to dimly fluorescent biological molecules. To obtain an accurate FLIM measurement, the percentage of “background photons” should be 1% or less of the total photons recorded. This can be achieved by making use of FLIM biosensors which are sufficiently bright in both their unbound state and their bound state, such as G-Ca-FLITS, and ensuring that sufficient protein is produced in the biological system of choice. According to our experience, a photon budget of 50,000 photons detected per region of interest (such as a cell or a subcellular region) results in a robust lifetime measurement through decay fitting, with a χ^2^ of 1 ± 0.2 and a high signal to background ratio.

Moreover, during imaging of live cells, it is necessary to balance several factors simultaneously. First, to increase the number of photons detected and thus reach the desired value of photons detected per region of interest, there are two options: increasing the laser power and decreasing the detector’s scan speed. However, both of these options negatively affect other aspects of the experiment: higher laser intensities necessarily result in a higher rate of phototoxicity and photobleaching of the fluorescent biosensor, while slower scan speeds increase the frame acquisition time, thus reducing time resolution in timelapse imaging. The latter can be especially problematic in the case of Ca^2+^-biosensors, since certain biological processes induce Ca^2+^ fluxes with changes in concentration that take place on the sub second scale (e.g., cardiac calcium sparks and neuronal calcium transients). Time resolution is also affected by spatial resolution as both higher pixel density (i.e. a smaller pixel size and thus greater spatial resolution) and a greater number of z-steps increase image acquisition time, thus reducing time resolution.

A final consideration in Time-Correlated Single Photon Counting FLIM is the frequency of the pulsed laser. Because of the relatively long lifetimes of new generation Ca^2+^ biosensors, such as G-Ca-FLITS (>4 ns in the Ca^2+^-free state), high-frequency pulsed lasers can lead to a photon pile-up effect and artificially bias the measurements towards shorter lifetimes. A 40 MHz frequency (25.6 ns measurement window per pulse) is preferable. Similar consideration should be taken to keep the number of photons detected per pixel per laser pulse below 1, since going above this number can similarly result in photon pile-up. (Fig. 2).

**Figure 2:**
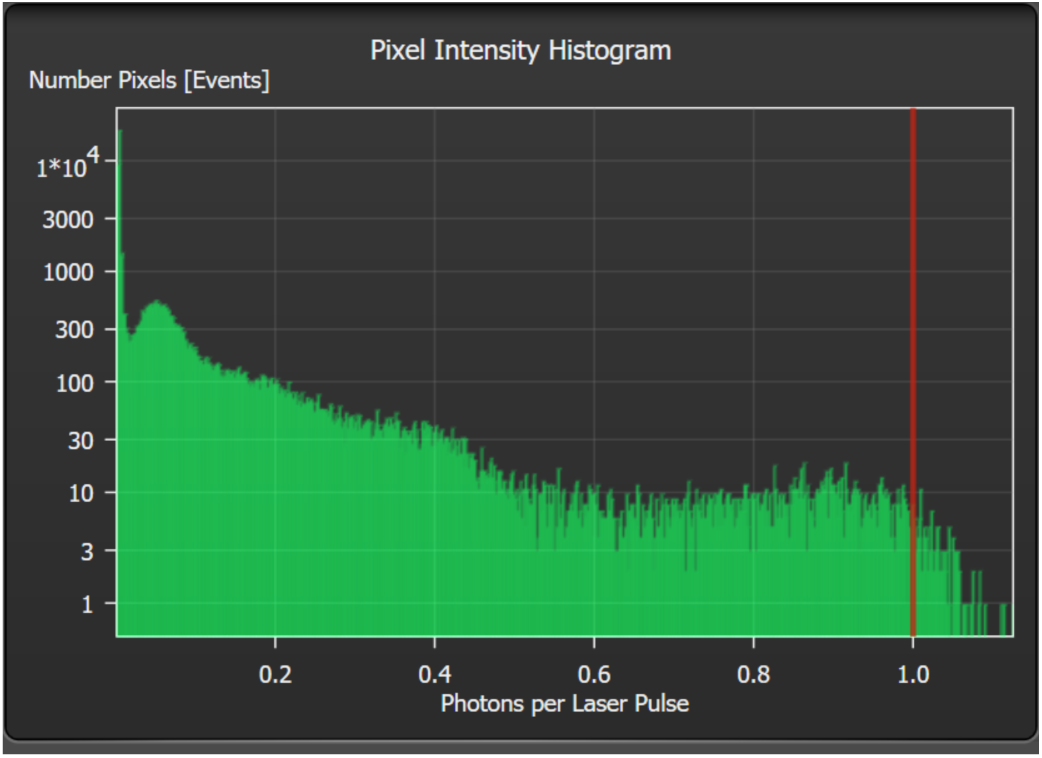
Screenshot of the diagram that shows the number of photons per laser pulse on LAS X. Care should be taken to collect most of the events at a rate of 1 photon (indicated by the red line) or less to avoid photon pile-up.

We can thus provide our imaging settings, which are optimized for the purpose of imaging G-Ca-FLITS-expressing adherent cells (thus a 2D experimental setup) on a 40x objective (HC PL APO 40x/0.95 NA DRY objective) on a Leica Stellaris confocal microscope.

- Frame resolution of 512 x 512, line accumulation at 2 (Note 24) and 200 Hz scanning speed: together these settings result in a pixel size of ∼500 nm and a frame time of 5 seconds, which fit our experiments as we measure the lifetime of individual whole cells and we image long-lasting (> 1 second) Ca^2+^ fluxes.
- Pinhole size of 1.5 AU at 530 nm (122.5 µm): since we are imaging whole cells, we open the pinhole enough to detect all photons originating from the entire cell at once (Note 25).
- White light laser at 85% power and 40 MHz frequency, 488 nm laser line at 7.5% power: This corresponds to a light at sample value of 2246.18 W/cm2 on our microscope, with which we can reliably reach the 100,000 photons threshold.
- HyDx in photon counting mode with detection range of 495-575 nm: When imaging a single construct we maximize signal by setting the beginning wavelength of the detection range at 7 nm after the laser line in use (thus 488 nm+7 nm=495 nm), and the end of the detection range at 80 nm after (thus 495 nm+80 nm=575 nm).

These settings have worked well for our experiments, which mainly consist of timelapse imaging of fields of view containing multiple adherent cells undergoing [Ca^2+^] changes lasting longer than 1 s (Fig. 3). Other experimental setups may require adjustments depending on the biological phenomenon being imaged. For example, measurement of Ca^2+^ fluxes at the subcellular scale (e.g., mitochondrial Ca^2+^ fluxes) may benefit from an increase in spatial resolution, which can be achieved by decreasing pixel size through an increase in format or an increase in magnification/zoom. Moreover, imaging of three-dimensional systems such as organoids or brain slices may require higher z-axis resolution, thus decreasing the size of the pinhole could be beneficial. Similarly, imaging of sub second Ca^2+^ fluxes will require an increase in scan speed or a decrease in line accumulation to decrease frame acquisition time.

**Figure 3:**
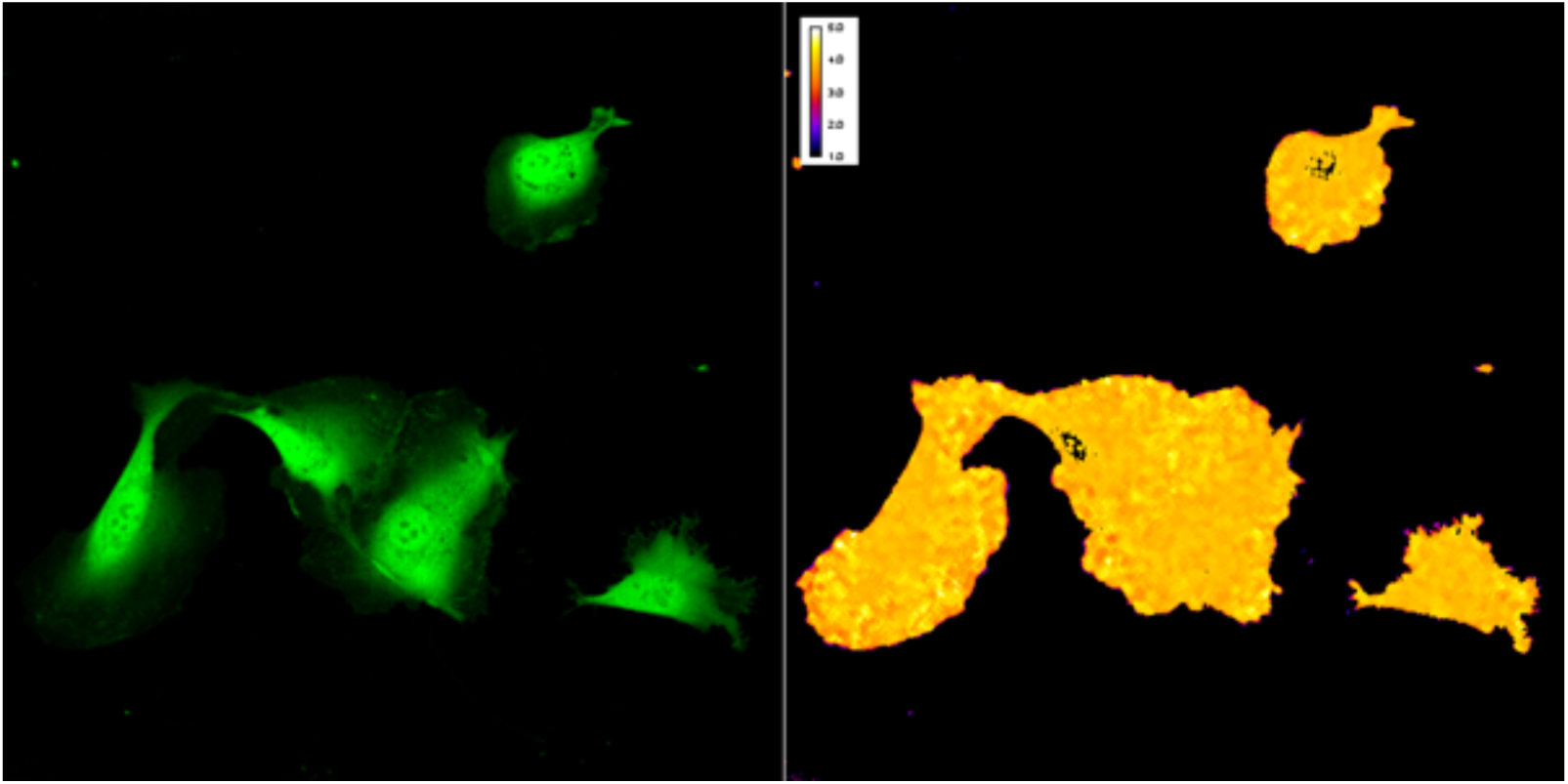
Confocal lifetime image of HUVEC expressing G-Ca-FLITS, 48 hours after electroporation. Left: Fluorescence intensity channel. Right: Lifetime channel (see Fig. 8).

### 3.8 FLIM Data analysis

FLIM data of live-cell experiments can be highly complex, because each pixel contains the arrival information of up to thousands of photons. As the dimensions of the image increase (which is the case for high-speed, high resolution Ca^2+^ imaging, or when multiplexing probes) the amount of information contained in a single FLIM timelapse increases accordingly. There are multiple approaches to analyze these large datasets, each with its own set of advantages and disadvantages. Example data is available from a zenodo archive ((DOI: 10.5281/zenodo.19357685)) and the code for visualizing the data is shared via a Github repo.

#### 1) Decay fitting

The FLIM module that is part of Leica Application Suite X (LAS X) allows fitting of the measured fluorescence decay curve to a theoretical decay model. We use the n-Exponential Reconvolution fit model, with 3 exponential components found from in vitro characterization of G-Ca-FLITS (Note 26). The mean 𝝉 is either Intensity or Amplitude weighted (Note 27). By drawing ROIs around individual cells it is possible to obtain a fitted mean lifetime (𝝉) per cell per acquired frame, which can be plotted to quantify the lifetime response of the sensor per cell over time (Fig. 5). Lifetime fitting is an accurate method for analyzing fluorescence lifetimes, however it is important to keep in mind that errors in the fit model can lead to biases in the data. The analysis of large amounts of data with LAS X can also be quite labor intensive.

The following is a step by step guide to go from our example raw live-cell FLIM data to a fitted-lifetime plot using LAS X for analysis and Python for plotting:

1. Open the “Timelapse-HUVEC.lif” project file on the LAS X FLIM module, and select the “TL_HUVEC_FLITS_Histamine” file (Alternatively, acquire live-cell lifetime data following section 3.7).
2. Click the “FLIM” button on the analysis method section (Fig. 4, A).
3. Draw ROIs around cells by using the options on the ROI panel (Fig. 4, B). Click the “New” button for each new ROI, otherwise they will be considered as the same ROI for lifetime fitting.
4. Once all ROIs are drawn, check the “Split” option on the FLIM settings panel (Fig. 4, C) to split the lifetime data into individual frames.
5. Click “Select Fit Model …” in the Fit panel (Fig. 4, D) and select n-Exponential Reconvolution.
6. Optional: Select the fitting range on the Decay curve (Fig. 4, E)/Fit panel to 0.5-20 ns (See Note 16 for a detailed explanation on the range chosen for G-Ca-FLITS).
7. If the lifetime components are not known, ensure all boxes on the “Fit” column (Fig. 4, F) within the Fit panel are checked, and beginning with 1 Exponential Component click “Fit All”. The Fit panel shows the obtained values for the currently selected row on the results table (Fig. 4, G). Check the χ^2^ values for ROI data (not for Overall decay, since this takes into account background) and if necessary, increase the number of components and fit all again until χ^2^ < 1.2 (Note 26). If the number and lifetime of the components are known (such as with G-Ca-FLITS), these can be typed into their respective rows in the Fit panel. Unchecking the “Fit” box forces the model to use the provided values, rather than calculating them from the data. Here, using the example data, we use three lifetimes, 0.76 ns, 2.07 ns, 4.56 ns that are derived from the fitting data of the purified biosensor.
8. For plotting the lifetime results using our custom python script, select all entries on the results table by clicking on a row and typing Ctrl + A.
9. Right click the selected results and select “Export Selected Rows…”, save as a .csv file. The data, which will be visualized in the next steps, is provided as “TL2_3componentFit.csv”.
10. Make sure that the conda environment is available (see step 3.2.14)
11. Launch Spyder and open the script: “Stellaris_LifetimePlotter_Simple.py”. After setting the appropriate file path, name, run the script by typing F5 or clicking the “Run File” button. Alternatively, from the conda environment, run the jupyter notebook “Stellaris_LifetimePlotter_Simple.ipynb”.
12. The result is a plot as displayed here in Fig. 5.

**Figure 4:**
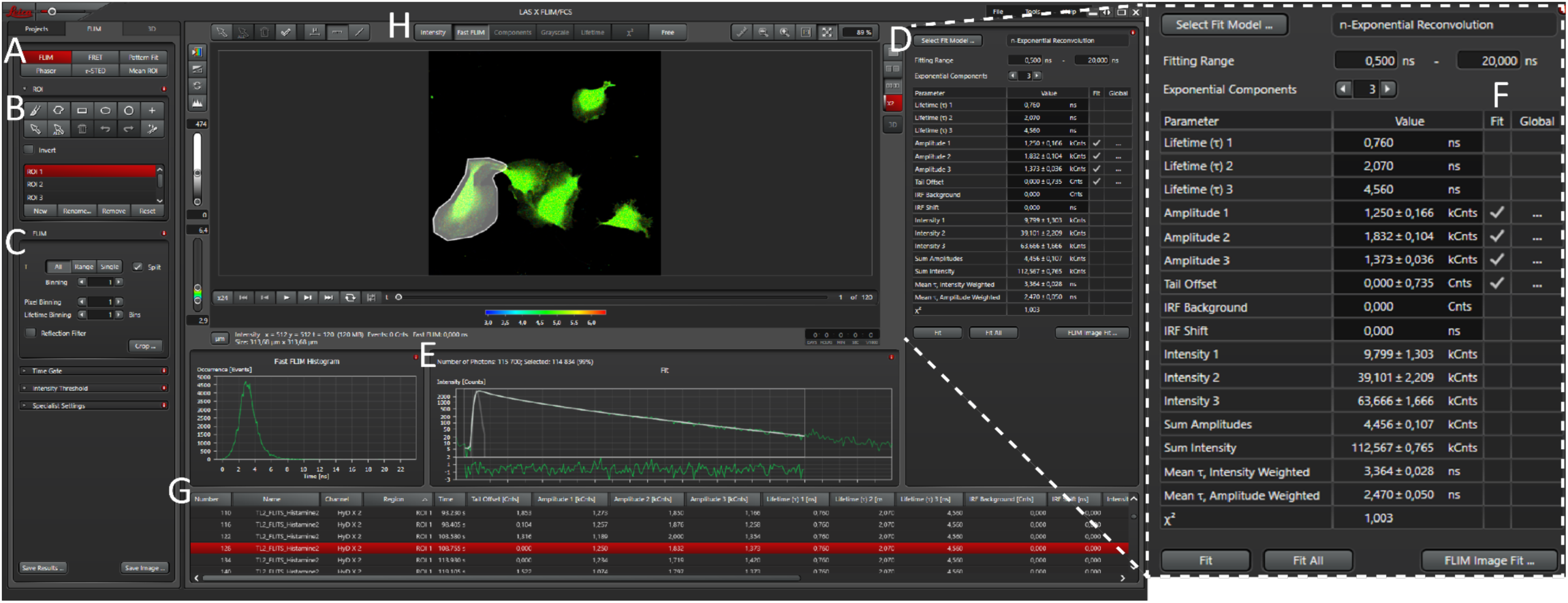
Screenshot of the LAS X FLIM software. Left: LAS X FLIM module. Right: Detail of the Fit panel. A-H indicates different components of the window. Decay curves were fitted per cell and per frame, using 3 exponential components with lifetime values obtained from in vitro characterization of G-Ca-FLITS.

**Figure 5:**
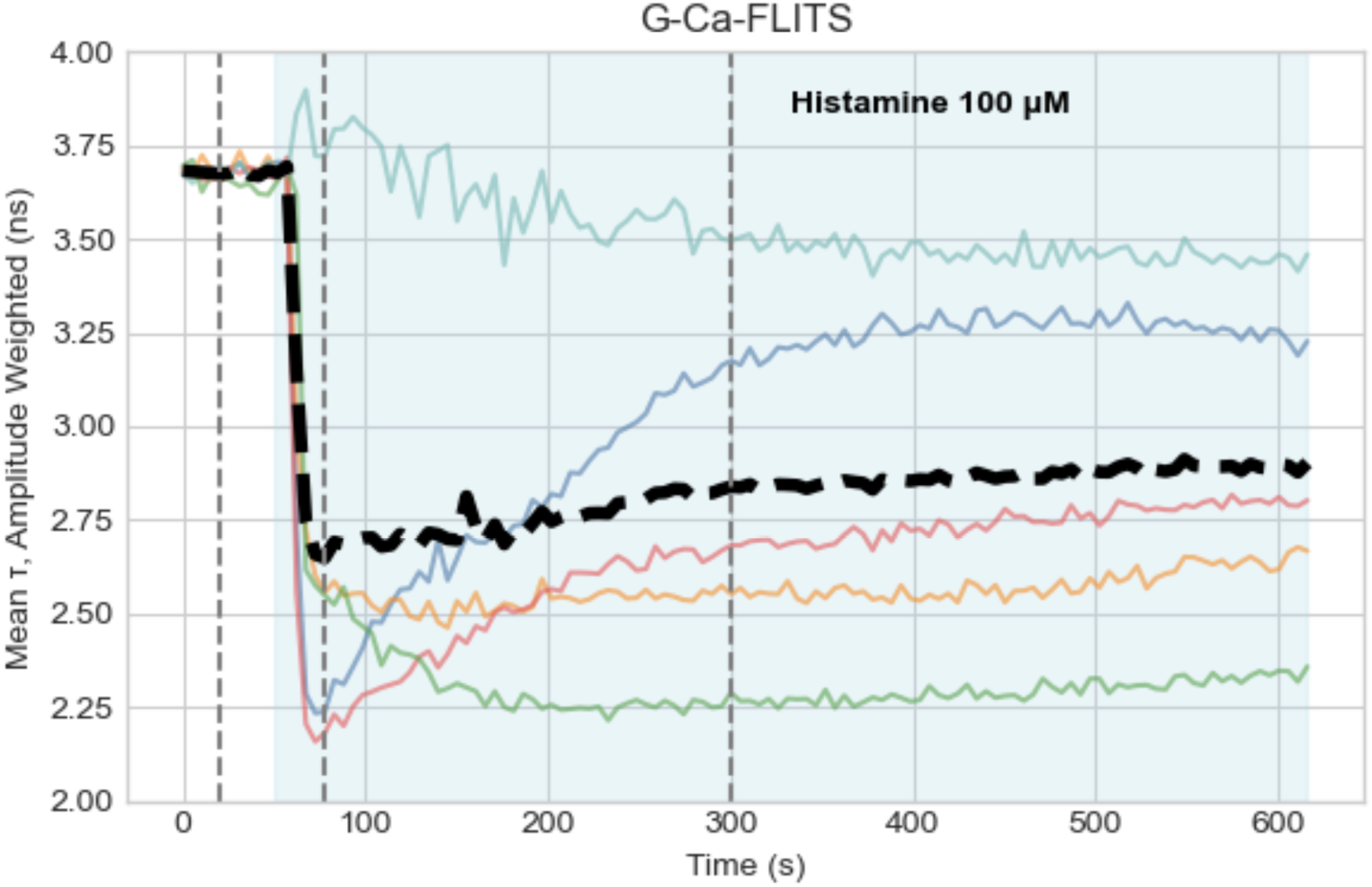
Plot showing the Amplitude Weighted mean 𝝉 (ns) of multiple HUVEC expressing G-Ca-FLITS (same cells shown on Fig. 3). The increase in intracellular Ca^2+^ is observed as a decrease in the lifetime of the biosensor after 100 µM histamine stimulation (shaded area). Colored lines represent individual cells, and the bold dashed line shows the mean response. Vertical dashed lines indicate timepoints shown as phasor plots and mean photon arrival times on Fig. 6 and Fig. 8, respectively.

#### 2) Phasor analysis

Phasor analysis is done by transforming the lifetime decay data contained in each pixel by using Fourier space and giving it two coordinates (G and S). With these coordinates it is possible to represent each pixel from an image as a point on a plot that corresponds to a given lifetime [25]. The coordinates on the phasor plot where points cluster are interpreted as bias-free average lifetimes. Lifetimes that land on the semicircle are considered monoexponential, while anything that falls within the circle is a mix of two or more lifetime components. Besides its low bias, an advantage of phasor analysis is that it is relatively straightforward to perform (possible on the LAS X software, although alternative open-source pipelines exist).

Step by step for obtaining phasors of live cell data on LAS X:

1. With your file open, click the “Phasor” button on the analysis method section (Fig. 6, A).
2. Check the “Split” option on the Dimensions panel (Fig. 6, B). This option displays an individual phasor plot per acquired frame, rather than an accumulation of the data from the entire timelapse. Drag the bar below the FLIM image (Fig. 6, C) to change the displayed frame and phasor plot.
3. Select the desired Phasor filter and threshold option (Fig. 6, D): A median filter of 15 and threshold of 5 were used to generate the plots on Fig. 6. Please refer to the LAS X user manual for details on the different filter options.
4. By default, the Phasor viewer already contains a circular lifetime cursor (Fig. 6, E), more can be added by clicking the “Draw cursor” button (Fig. 6, F) and then clicking anywhere in the plot.
5. To obtain a lifetime measurement, click and drag the circular cursors so that the clustered data points on the plot fall within their radius (the radius can be modified). The lifetime for the position of each cursor is displayed on the “Data display of the components panel” (Fig. 6, G). Furthermore, the pixels on the image with lifetimes that fall within the cursor are displayed with the corresponding color on the FLIM image.
6. Phasor plots for individual cells can be obtained by drawing ROIs and selecting them on the Display of the loaded image data records (Fig. 6, H).
7. Phasor plots can be exported as 3-channel RGB images (either as animated timelapses or single accumulated plots, depending on whether the Split checkbox is active) by selecting “Save Phasor …” (Fig. 6, I). Alternatively, right click on the phasor plot and choose “Export Image …”.

**Figure 6:**
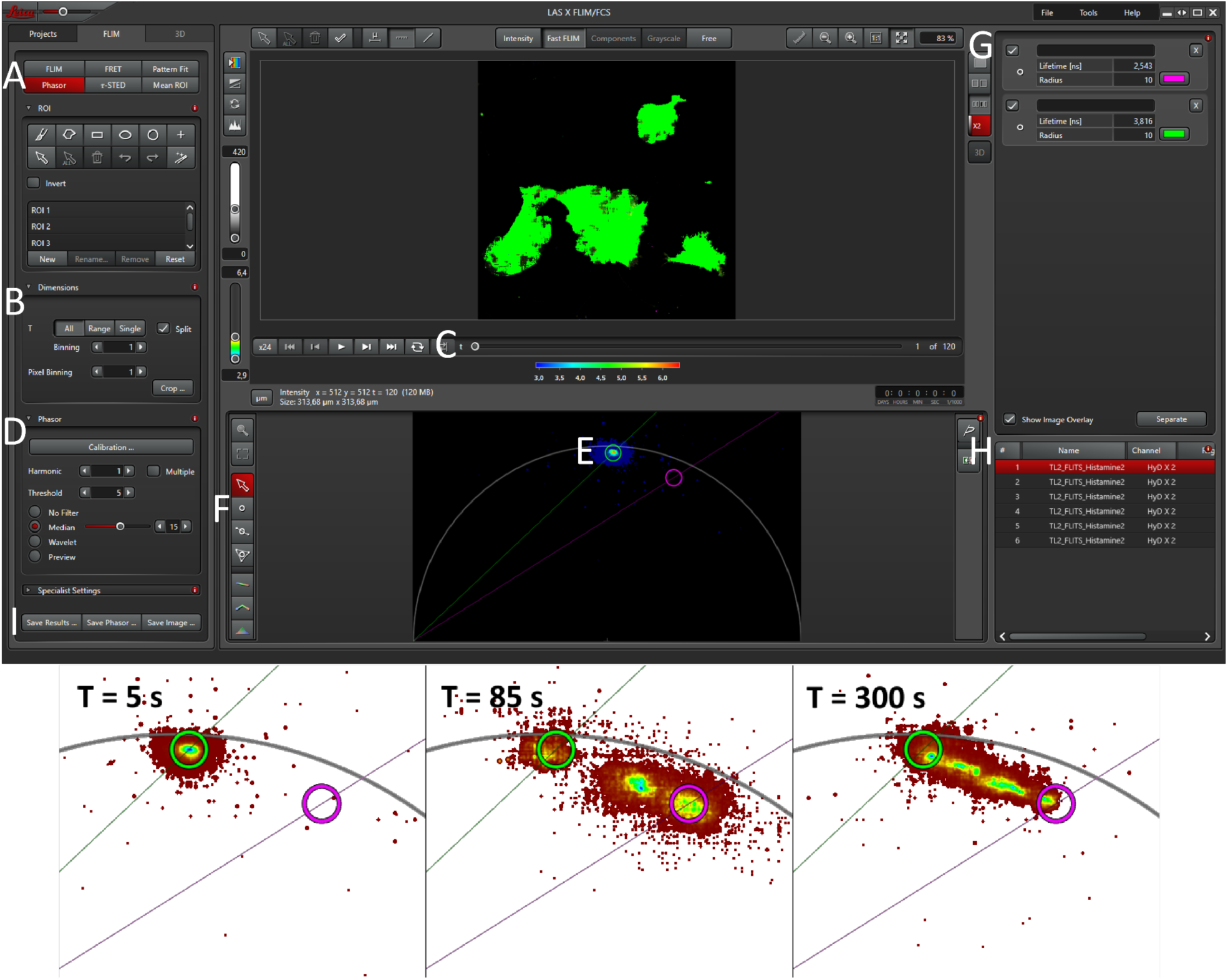
Phasor analysis of FLIM data. Top: Phasor window in the LAS X FLIM module, A-I indicates different components of the window. Bottom: Phasor plots showing the lifetime of G-Ca-FLITS in HUVEC before (T= 5 s), at peak response (T= 85 s), and during recovery (T = 300 s) following 100 µM histamine stimulation. Circles represent basal (green, 3.82 ns) and peak stimulation (magenta, 2.54 ns) G-Ca-FLITS lifetimes. Individual cells, or groups of cells with similar lifetimes can be observed as clusters of points in green-blue colors.

#### 3) Mean photon arrival time (FastFLIM on LAS X)

A fast way of visualizing lifetimes directly on the acquired image is by calculating the average photon arrival time in each pixel. This method reduces the complexity of the information contained in each pixel to a single mean value, which can be easily color-coded and displayed as a LUT. The mean photon arrival times are considered less accurate since they are not fitted to a model that takes into account the number or contribution of each lifetime component. It can be, in any case, a convenient method for exploring the spatiotemporal dynamics of a biosensor response. Our group routinely uses a custom Fiji macro written for this purpose:

1. On LAS X: Ensure “FastFLIM” is selected on the FLIM image viewer (Fig. 4, H), right click the image and select “Export raw image”.
2. On the LAS X Save Image window, choose the following options (Fig. 7, left):

○ ImageJ TIFF
○ Fixed Range
○ Intensity [Cnts] | per Grey Level | 1.000
○ Lifetime (t) [ns] | Range | 0.000-10.000
3. The export should result in a 2-channel TIFF file: Channel 1 contains the intensity data and channel 2 the mean photon arrival times per pixel. The exported data is available in the zenodo archive as “TL_HUVEC_FLITS_Histamine.tif”.
4. Open Fiji and open the file “TL_HUVEC_SLITS_Histamine.tif” by dragging it into the Fiji window.
5. Open the macro “Display_tau_Stellaris_v5_Stacks.ijm” by dragging it into the Fiji window.
6. Click “Run” on the macro edition window that opens, or type Ctrl + R.
7. Set parameters on the “Input for Lifetime Display” window. For the data on Fig. 8, the following relevant settings were used (Fig. 7, right). Settings need to be adjusted based on the data:

○ What to process: Process current image
○ Minimal Tau for display (ns): 1
○ Maximal Tau for display (ns): 5
○ Automatic determination of Intensity threshold: No
○ Minimum image intensity for lifetime statistics/display [Cnts]: 10
○ Maximum image intensity for lifetime statistics/display [Cnts]: 1000
○ Median filter size to smooth lifetime image: 3
8. Click “OK” on the “Input for Lifetime Display” window.
9. The resulting stack can be saved by using the “Save As” option under the File menu on the Fiji window.
10. Mean photon arrival times per frame are temporarily stored in the Fiji Log window, and can be saved as a .csv file from there.

**Figure 7:**
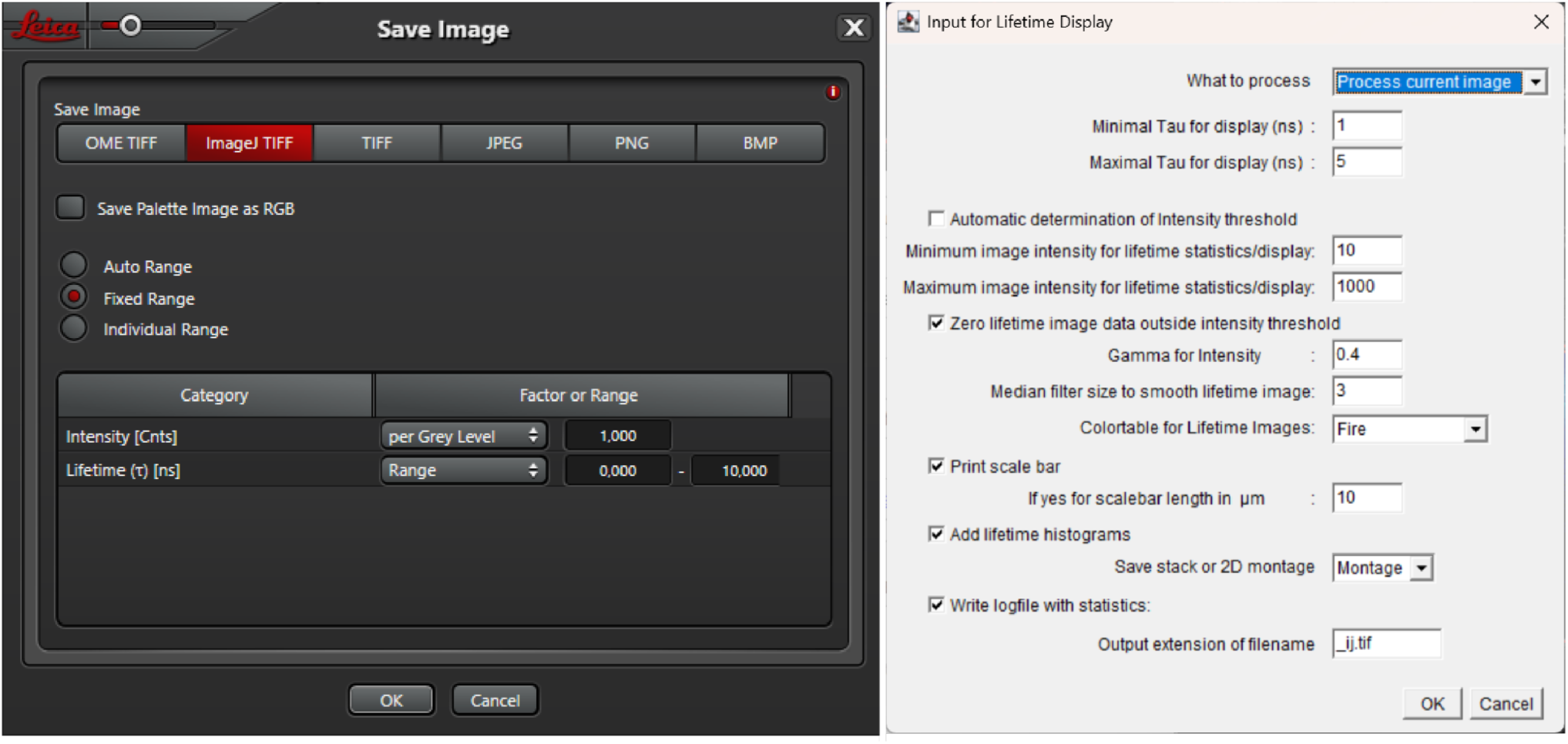
Screenshots of software for processing the mean arrival time data. Left: Settings for exporting raw FLIM data from LAS X for analysis with the FIJI macro. Right: “Display_tau_Stellaris_v5_Stacks.ijm” parameters used for Fig. 8.

**Figure 8:**
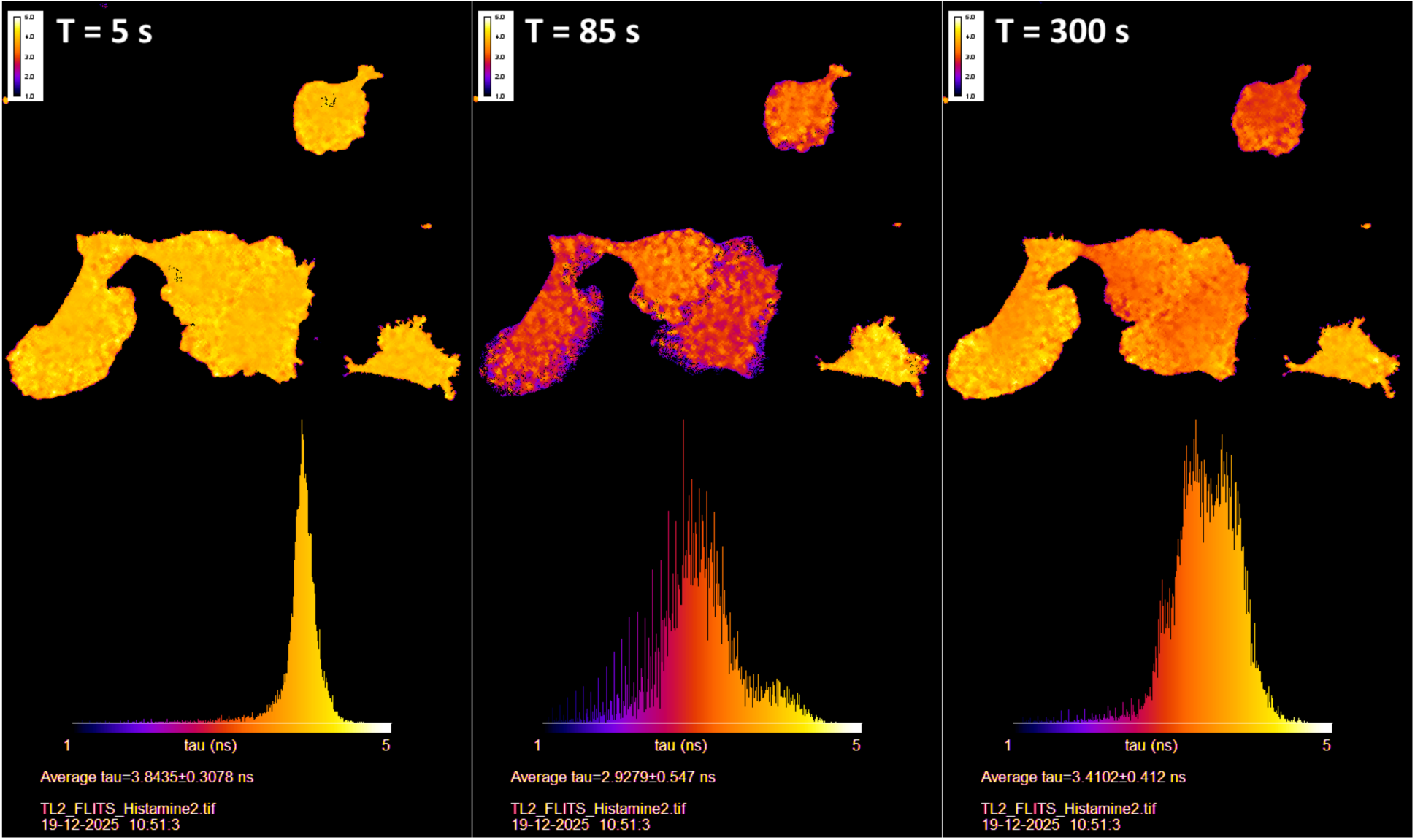
Average photon arrival times of G-Ca-FLITS in HUVEC before (T = 5 s), at peak response (T = 85 s), and during recovery (T = 300 s) following 100 µM histamine stimulation. Fake color LUT represents photon arrival time per pixel, with warmer colors indicating long lifetimes (low Ca^2+^) and colder colors indicating short lifetimes (high Ca^2+^). Histograms show the distribution of photon arrival times and an average lifetime for each image.

### 3.9 Conversion of FLIM data to [Ca^2+^] in live cells

An absolute quantification of Ca^2+^ based on FLIM measurements requires both a high-quality calibration dataset (i.e. imaging of a FLIM sensor at known [Ca^2+^], as outlined in 3.2 steps 1 to 11) and high-quality experimental dataset (i.e. acquired as outlined in 3.7). Moreover the imaging conditions (i.e. ambient pH and temperature) and the imaging settings (especially with regard to laser line used, laser pulse frequency and detection spectral range) should be the same for both datasets, and the lifetime should be estimated in the same way for both datasets, whether that be through fitting of the fluorescence decay curve or fitfree approaches. This degree of similarity is necessary for the results of the quantification to be accurate. A robust and straightforward approach is to make use of the LAS X software’s FLIM module to determine G-Ca-FLITS’ lifetime in both the calibration and experimental datasets, and convert the experimental dataset’s lifetimes into [Ca^2+^] using an R-based pipeline.

1. Acquire an in vitro calibration dataset by imaging purified sensor samples at known [Ca^2+^] as outlined in protocol 3.2 steps 1 to 11.
2. In the LAS X software, fit the decay curves of the in vitro calibration dataset as outlined in protocol 3.7 in order to establish the individual lifetime components that make up the fluorescence decay of the samples. In the case of G-Ca-FLITS we obtained an acceptable fit (χ^2^ <1.2 and low residuals) with a three components fit: 𝝉1 = 0.76 ns, 𝝉2 = 2.07 ns, 𝝉3 = 4.56 ns.
3. Export the data as a .csv file. The data for the calibration is available as “Calibration_fit.csv” in the Github repo.
4. Image cells expressing the sensor using the same imaging settings and at the same temperature and pH as those of the in vitro calibration dataset.
5. In the LAS X Software, draw any cellular or subcellular ROIs as wanted.
6. Use the component lifetimes determined by fitting the in vitro calibration dataset’s decays to fit the fluorescence decay curves of the cell-based experimental dataset.
7. Export the fitting results, saving the table as a .csv file. The data for the example timelapse is available in the github repo as “Experiment_Fit.csv”.

In order to convert the lifetime information of the experimental dataset to Ca^2+^ concentrations, the fit data is processed through an R-based pipeline. The scripts as well as the example calibration and experimental datasets relevant to this protocol can be found in the folder “3.9_Converting_FLIM_data_to_[Ca2+]_in_live_cells” of the github repository MMB_Quantitative_imaging_of_calcium_dynamics. First, similarly to what is done in protocol 3.2 using line fraction values, the intensity weighted mean 𝝉 values calculated during the fitting of the calibration dataset are used to establish the constants in the equation which dictates the relationship between the intensity weighted mean 𝝉 value of each sample and its [Ca^2+^] (Equation 3).

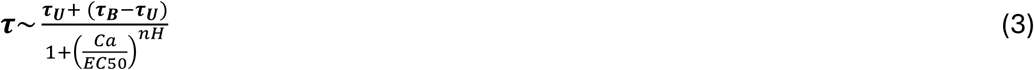

where 𝝉 is the intensity weighted mean 𝝉 value; 𝝉_𝑼_ is the maximum 𝝉 value, corresponding to the 𝝉 of the 0 [Ca^2+^] sample within the calibration dataset in which G-Ca-FLITS’ Ca^2+^-unbound state is most abundant; 𝝉_𝑩_is the minimum 𝝉 value, corresponding to the 𝝉 of the 39 μM [Ca^2+^] sample within the calibration dataset in which G-Ca-FLITS’ Ca^2+^-bound state is most abundant; *Ca* is the [Ca^2+^] that corresponds to the 𝝉 value; *EC50* is the [Ca^2+^] in which G-Ca-FLITS’ lifetime corresponds to the middle point between the lifetime measured in the 0 [Ca^2+^] sample and the 39 μM [Ca^2+^] sample; *nH* is the Hill coefficient of the sensor’s Ca^2+^-induced lifetime change.

Once the equation’s constants are determined, they are used to calculate the [Ca^2+^] value corresponding to each intensity weighted mean 𝝉 value in the experimental dataset (Equation 4).

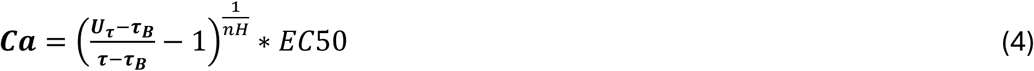

where 𝝉 is the intensity weighted mean 𝝉 value; 𝝉_𝑼_ is the maximum 𝝉 value, corresponding to the 𝝉 of the 0 [Ca^2+^] sample within the calibration dataset in which G-Ca-FLITS’ Ca^2+^-unbound state is most abundant; 𝝉_𝑩_ is the minimum 𝝉 value, corresponding to the 𝝉 of the 39 μM [Ca^2+^] sample within the calibration dataset in which G-Ca-FLITS’ Ca^2+^-bound state is most abundant; *Ca* is the [Ca^2+^] that corresponds to the 𝝉 value; *EC50* is the [Ca^2+^] in which G-Ca-FLITS’ lifetime corresponds to the middle point between the lifetime measured in the 0 [Ca^2+^] sample and the 39 μM [Ca^2+^] sample; *nH* is the Hill coefficient of the sensor’s Ca^2+^-induced lifetime change.

8. Download the github repository “MMB_Quantitative_imaging_of_calcium_dynamics” from (https://github.com/AndreaCaldarola/MMB_Quantitative_imaging_of_calcium_dynamics) as a zip and extract the folder “3.9_Converting_FLIM_data_to_[Ca2+]_in_live_cells” to an appropriate location on your computer.
9. Move the .csv files containing the fitting results as well as a .csv file containing the information on the [Ca^2+^] of each sample of the in vitro calibration dataset (as in steps 10 and 11 of protocol 3.2) into the extracted folder.
10.In RStudio, open the file “Mean_Lifetime_intensity_Weighted_to_Ca.qmd” located within the extracted folder.
11. Install all package dependencies when prompted.
12. If not using the provided G-Ca-FLITS sample datasets, change the file names within the “read.csv()” lines of code to those of your files.
13. Hold down the Ctrl, Alt and R keys on the keyboard (option+cmd+R on a Mac) in order to fully execute the script.
14. A new folder called “Plots” should be created, containing plots which showcase the fit-based sensor calibration and the changes in [Ca^2+^] that took place during the cell-based timelapse imaging experiment (Fig. 9).

## 4. Expected outcomes

Figure 9 is a summary of the expected outcome, showing the calibration curve derived from the purified protein and the plots that report on the quantitative Ca^2+^ measurements in HeLa cells and how these change dynamically in response to histamine.

Basal Ca^2+^ concentrations are below 59 nM, which is the lower limit of detection for this probe. Upon histamine addition, the Ca^2+^ concentration rises rapidly to micromolair concentrations. Subsequent Ca^2+^ transients rise up to 1 µM. Both the images (fig. 9F) as well as the plots (figures 9C and 9D) show substantial heterogeneity between cells. These results are in line with previous observations [14, 21]

**Figure 9:**
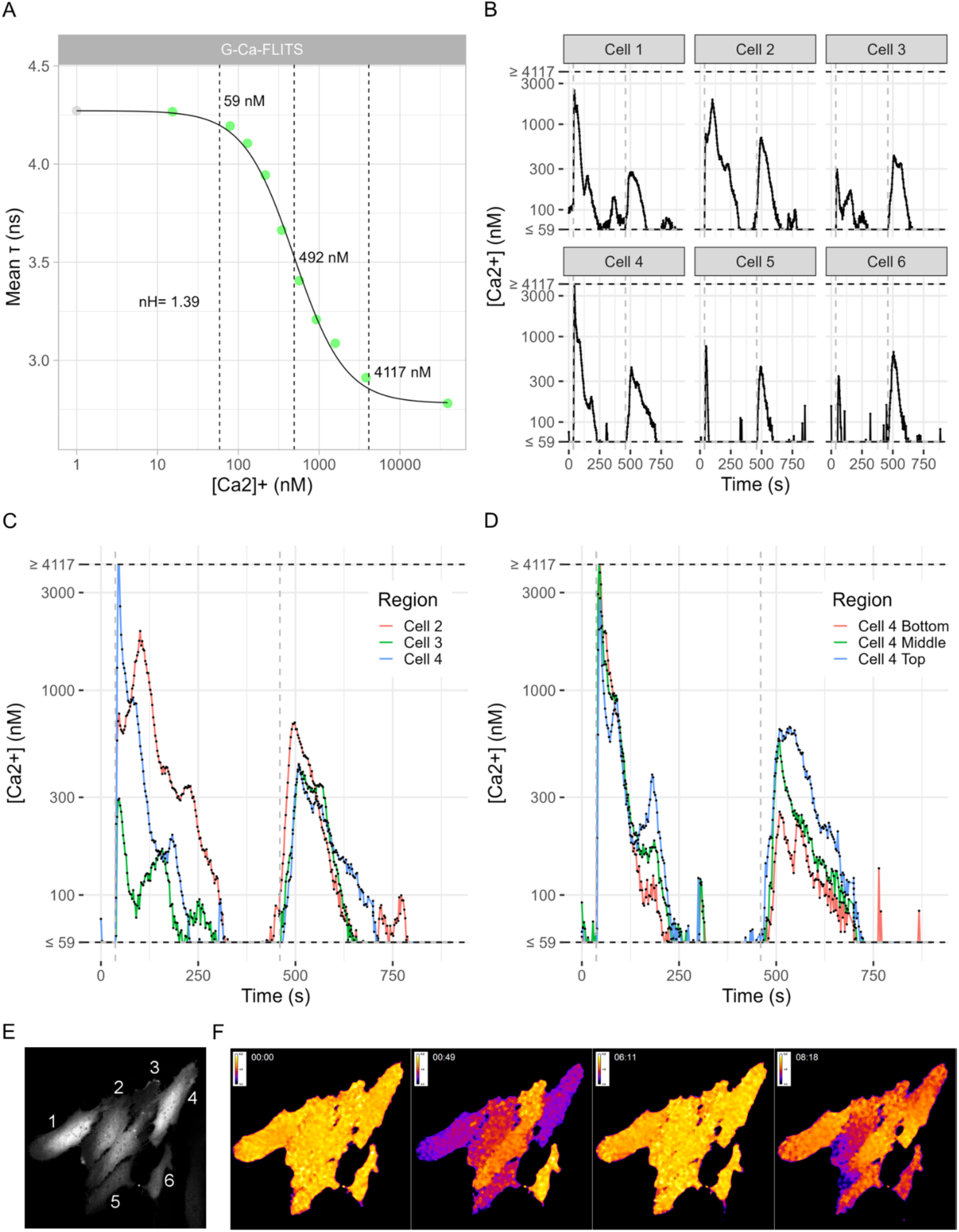
Converting FLIM data to [Ca^2+^] in live cells. A) Fitted mean 𝝉 plot: the mean 𝝉 of each of the calibration dataset samples is plotted as a function of its [Ca2+]. The 0 [Ca2+] sample (shown in gray) is plotted as 1 nM [Ca2+] in order to display it on a logarithmic axis. The Hill coefficient (nH), EC05, EC50 and EC95 values are shown, as well as a black line of points computed using the fitting results. B, C & D) Plots showing the changes in [Ca^2+^] that took place during the sample cell-based G-Ca-FLITS imaging experiment dataset. In all plots two horizontal dashed lines are present, which correspond to the EC05 and EC95 [Ca^2+^] of G-Ca-FLITS’ Ca^2+^-induced lifetime change. We limit our [Ca^2+^] quantification to lifetimes that correspond to Ca^2+^ values within this range, as this is the range in which G-Ca-FLITS is most responsive to changes in [Ca^2+^]. Two vertical gray dashed lines are also present, which indicate the time points in which cells were treated with 100 μM Histamine (left-most dashed line, t= 37 s) and 2 μM Thapsigargin (right-most dashed line, t = 460 s). E) First frame of the cell-based timelapse experiment represented in plots B, C and D. The intensity channel is shown and each cell is numbered according to its name in the plots. F) Four frames of the cell-based timelapse experiment represented in plots B, C and D. The cells’ average photon arrival times are shown before any stimulation (frame 1, t = 0 s), at the peak of the cells’ response to 100 μM Histamine (frame 2, t = 49 s), having mostly returned to basal [Ca^2+^] after the Histamine response (frame 3, t = 371 s) and finally at the peak of the cells’ response to 2 μM Thapsigargin (frame 4, t = 500 s).

## 5. Notes

**Note 1**: When only washing cells without trypsinization, PBS should be supplemented with 1 mM CaCl2 and 0.5 mM MgCl2 (PBS++) to avoid breaking cell junctions.

**Note 2**: For live-cell imaging, cells are plated on round coverslips (diameter 24 mm and thickness #1.5) placed in a standard 6-well plate. The coverslips are then placed and imaged in imaging cell chambers (Such as ThermoFisher #A7816). Alternatively, cells are plated and imaged directly on 12 or 24 well glass-bottom plates (Mattek corp.).

**Note 3**: Ideally, the DNA concentration of plasmids for transfection is >100 ng/μL, with a A260/A280 purity of >1.80.

**Note 4**: A 100 mm petri dish at ∼90% confluency is enough for 4 *n* electroporations. Use ideally 250,000-500,000 cells per electroporation *n*.

**Note 5**: Endothelial cells are very sensitive to the presence of bacterial endotoxin in purified DNA. For best results, DNA purification for endothelial microporation should be done with a mini/maxiprep protocol that includes endotoxin-binding reagent as a purification step (e.g., ThermoFisher #K0861).

**Note 6**: This version of the electroporation kit has been discontinued in favor of a newer version, untested in our lab. Reagents and equipment (tips, tubes) for the kit can be regenerated, see [26]).

**Note 7**: We image HeLa and most cell lines in a low-background Microscopy Medium (20mM HEPES pH=7.4, 137 mM NaCl, 5.4 mM KCl, 1.8 mM CaCl_2_, 0.8 MgCl_2_, and 20 mM Glucose in MQ water). Endothelial cells are imaged in their full growth EGM-2 medium.

**Note 8**: At the end of the 6 hour room temperature growth period, in the case of G-Ca-FLITS, the culture should develop a noticeable yellow-green color.

**Note 9**: Once the bacteria are frozen, it is possible to temporarily store them in this condition. In our experience the yield of the purification process is high, and storing the bacteria frozen for a week does not significantly affect the final yield.

**Note 10**: Freezing the bacteria horizontally enables faster defrosting in the next step.

**Note 11**: Due to the high centrifugation speed used, we deem it necessary to weigh the samples to be centrifuged on a standard laboratory scale and to ensure that all samples (and any tube used to balance the centrifuge) have equal weight. Small differences in weight can be corrected by adding ST buffer to the lighter samples.

**Note 12**: The elution of the purified sensor is not equal: the initial and final 33% of the eluted volume contain a lower concentration of the purified protein, while the intermediate eluate contains a higher concentration of the purified protein. Since in our case this can be easily determined by observing the color of the eluate, we simply gather the first, intermediate and final fraction in three different tubes, switching from the first to second tube once the color of the eluate becomes more intense, and from the second to the third once the color begins to fade. We only use the brightest, most concentrated fraction for the characterization of the sensor’s lifetime response.

**Note 13**: In the calibration kit we are using, the final free [Ca^2+^] of each sample is determined by the total [Ca^2+^] present in the sample (between 0-10 mM), and by the samples temperature and pH (which we do not change from the 7.2 pH of the provided buffers), as the Kd of the Ca^2+^ chelator used in the kit’s buffers, EGTA, is both temperature and pH-dependent.

**Note 14**: In order to produce a robust calibration curve, the free [Ca^2+^] of the sensor samples should reside within the range of [Ca^2+^] in which the greatest lifetime contrast can be observed. To construct our calibration curves we image at least 10 samples, and we aim for most of the samples to be to be evenly spread within the range of [Ca^2+^] which determine 5% and 95% of the sensor’s total change in lifetime when comparing the lifetime of the 10 mM K_2_EGTA (i.e. the sample in which we have the highest fraction of unbound sensor) and the 10mM CaEGTA sample (i.e. the sample in which we have the highest fraction of Ca^2+^-saturated sensor). This is the range in which the sensor is most responsive to changes in [Ca^2+^]. In order to find this range we recommend assembling a first “exploratory” calibration, with [Ca^2+^] centered around the Kd of the sensor’s binding domain, if known, or around a [Ca^2+^] estimated to be close to the sensor’s Kd based on exploratory cell-based imaging data.

**Note 15**: The Ca^2+^ buffering system used by us is sensitive to both pH and temperature, which must be taken into account when executing the calibration. Similarly, the photophysical properties of a sensor may be affected by temperature or pH. For this reason we highly recommend imaging the calibration samples at the same temperature and pH of the system that you ultimately want to use the sensor in. G-Ca-FLITS’s lifetime is known to be stable at pH above 7, thus we are confident that the results of a calibration acquired at pH 7.2 (which is the pH of the 0 and 10mM CaEGTA buffers provided in the kit used by us) can be used to accurately determine the [Ca^2+^] of cells at pH 7.4. We are however aware that both the lifetime and Kd of G-Ca-FLITS are affected by temperature, exhibiting a reduction in lifetime and increase in affinity when comparing measurements taken at room temperature and 37 °C. For this reason temperature control should be used both when imaging the sensor calibration samples and during cell-based imaging experiments.

**Note 16**: Following the python script execution the 1-dimensional arrays are saved as .csv files, namely “all_decays.csv” and “all_decays_raw.csv”. These arrays can be directly plotted within a spreadsheet editing software in order to obtain each sample’s decay profile, allowing for visual inspection of the raw data. In our experimental setup we detect a peak of photons in the 20-22 ns detection range, which can be observed by plotting the arrays in the “all_decays_raw.csv” file. This peak of signal does not originate from G-Ca-FLITS but is instead, to our estimation, caused by the detection of a reflection of the excitation laser. For this reason we “trim” the calibration samples’ arrays, removing all photons detected after bin 200, which corresponds to the first ∼19.5 ns of the 25.63 ns detection range. These trimmed arrays can similarly be observed by plotting the arrays present in the “all_decays.csv” file. In our longest lifetime sample (G-Ca-FLITS in absence of Ca^2+^), the photons detected in the 19.5 to 25.63 ns range amount to ∼2% of the total photons detected. Given that, we deem the loss of information due to the trimming of the last fraction of the fluorescence originating from G-Ca-FLITS as acceptable in order to avoid biasing all samples towards higher lifetimes (which would artificially decrease the sensor’s contrast) while maintaining a fit and assumptions-free approach.

**Note 17**: HUVEC are very sensitive in general, they should be kept out of the incubator as briefly as possible and not be pipetted too roughly.

**Note 18**: EGM-2 degrades quickly. Do not warm up for longer than 10 minutes before culturing cells, and store in the dark.

**Note 19**: If overexpression of the construct of interest is a concern,100 ng of the DNA of interest can be used, and the remaining 400 ng can be filled with non-coding, empty DNA vector. This maintains the most efficient transfection ratio while decreasing the total amount of coding DNA that is delivered to the cells.

**Note 20**: Gently pipetting at each step makes electroporation more consistent, since cells will settle down rather quickly.

**Note 21**: This is the most common cause for errors during electroporation. Bubbles in the pipette tip will prevent the electrical circuit from closing properly, which can lead to either a failed transfection or low cell viability.

**Note 22**: A short burst of bubbles but no electric arc should be visible near the electrode during the pulse. If electroporation fails because the circuit is not properly closed, ensure everything is firmly in place and that there are no bubbles inside the pipette tip.

**Note 23**: Electroporation naturally can kill about 30% of cells. Since HUVEC are quite sensitive to the presence of dead cells this step is important. An additional wash step with PBS++ may be necessary if too many dead cells are present.

**Note 24**: We prefer line as opposed to frame accumulation, as we want to prioritize having the smallest time interval between each repetition, due to the fast nature of Ca^2+^ fluxes. Line repetitions are dispensable but may help boost the photon count when needed, at the expense of time resolution/phototoxicity.

**Note 25**: When imaging thicker cell types, the pinhole size can be increased up to 3 AU.

**Note 26**: The number of components for a given lifetime decay depends on the characteristics of the sample. The χ^2^ value indicates how well the measured lifetime decay fits the number of components chosen. χ^2^ = 1 corresponds to a perfect fit, while values below 1 and above 1 indicate under- and overfitting respectively. Ideally, one should see a large decrease (>3-5 fold) in the value of χ^2^ when increasing the number of components from a number that is not sufficient to describe the lifetime decay (e.g., the χ^2^ obtained when fitting the fluorescence decay of any single FP lifetime-based biosensor at its EC50 [Ca^2+^] to a single component lifetime) to the “true” number of components which make up the measured lifetime. We consider χ^2^ values <1.2 as an indication that the fit describes the lifetime decay curve to an acceptable degree. Any small decrease beyond that as more components are added does not necessarily indicate a better fit.

**Note 27**: The value of 𝝉 may differ between the two weighing methods, we suggest using the one closest to theoretically expected lifetimes, or to lifetimes measured using low-bias methods (such as phasor analysis).

## Acknowledgments

We thank our colleagues at Molecular Cytology for their various contributions and support.

## Data availability

All data and code used in this manuscript is available:

Data: https://zenodo.org/records/19357685

Code: https://github.com/AndreaCaldarola/MMB_Quantitative_imaging_of_calcium_dynamics

